# Semaphorin 4B is an ADAM17-cleaved inhibitor of adipocyte thermogenesis

**DOI:** 10.1101/2022.10.11.511765

**Authors:** Abdulbasit Amin, Marina Badenes, Johanna Tüshaus, Érsika de Carvalho, Emma Burbridge, Pedro Faísca, Květa Trávníčková, André Barros, Stefania Carobbio, Pedro Domingos, Antonio Vidal-Puig, Luís Moita, Sarah Maguire, Kvido Stříšovský, Stefan F. Lichtenthaler, Colin Adrain

## Abstract

**Objective:** The metalloprotease ADAM17 (also called TACE) plays fundamental roles in homeostasis by shedding key signaling molecules from the cell surface. Although its importance for the immune system and epithelial tissues is well-documented, little is known about the role of ADAM17 in metabolic homeostasis. The purpose of this study was to determine the impact of ADAM17 expression, specifically in adipose tissues, on metabolic homeostasis.

**Methods:** We used histopathology, molecular, proteomic, transcriptomic, in vivo integrative physiological and ex vivo biochemical approaches to determine the impact of adipose tissue-specific deletion of ADAM17 upon adipocyte and whole organism metabolic physiology.

**Results:** ADAM17^adipoq-creΔ/Δ^ mice exhibited a hypermetabolic phenotype characterized by elevated energy consumption and increased levels of adipocyte thermogenic gene expression. On a high fat diet, these mice were more thermogenic, while exhibiting elevated expression levels of genes associated with lipid oxidation and lipolysis. This hypermetabolic phenotype protected mutant mice from obesogenic challenge, limiting weight gain, hepatosteatosis and insulin resistance. Activation of beta-adrenoceptors by the neurotransmitter norepinephrine, a key regulator of adipocyte physiology, triggered the shedding of ADAM17 substrates, and regulated ADAM17 expression at the mRNA and protein levels, hence identifying a functional connection between thermogenic licensing and the regulation of ADAM17. Proteomic studies identified Semaphorin 4B (SEMA4B), as a novel ADAM17-shed adipokine, whose expression is regulated by physiological thermogenic cues that acts to dampen thermogenic responses in adipocytes. Transcriptomic data showed that cleaved SEMA4B acts in an autocrine manner in brown adipocytes to dampen the expression of genes involved in thermogenesis, adipogenesis, lipid uptake, storage and catabolism.

**Conclusion:** Our findings identify a novel ADAM17-dependent axis, regulated by beta-adrenoceptors and mediated by the ADAM17-cleaved form of SEMA4B, that may act to limit uncontrolled energy depletion during thermogenesis.

## 1. Introduction

Energy balance, the net outcome of caloric intake versus caloric expenditure, is controlled by complex communication amongst metabolic organs (e.g. brain, nervous system, adipose tissues, liver, pancreas, immune system and skeletal muscles) [1,2]. As a consequence of the rising obesity epidemic in developed and middle-income countries, extensive research efforts are dedicated towards identifying pathways that can be drugged to promote weight loss in obese subjects [3–5]. Adipose tissues are organs that play central roles in energy balance, organismal homeostasis and are implicated in a range of metabolic and infectious diseases [6–10]. They are specialized into 2 major functional classes: white adipose tissue (WAT) and brown adipose tissue (BAT). In the WATs, adipocytes store excess calories as triglycerides in a single lipid droplet comprising about 90% of cell volume. In the BAT, adipocytes contain multiple lipid droplets, are enriched in mitochondria and are highly catabolic [7].

One of the key ways in which adipocytes communicate, locally within an adipose tissue, or hormonally to distant target organs, is via secreted signaling molecules collectively known as adipokines. Prominent examples of adipokines include fibroblast growth factor 21 (FGF21), a protein that regulates hepatic gluconeogenesis, and enhances insulin sensitivity and thermogenesis in adipose tissue [6,11]. Another adipokine, the hormone leptin regulates energy balance centrally via the control of satiety [12,13]. Adipose tissues are themselves subject to incoming hormonal signals. They are richly innervated by sympathetic nerve endings that secrete the neurotransmitter norepinephrine (NE) in response to physiologic stimuli such as cold, overnutrition, and fasting [14–19]. WAT serves as reservoir of energy that can be mobilized when required (e.g. in the postprandial state, fasting, or during infection). The cue for energy release from WAT is signaled by NE, an activator of β-adrenergic receptors [19]. This triggers lipolysis in lipid droplets: the breakdown of triglycerides into their constituent free fatty acids (FFAs) and glycerol, which are released into the systemic circulation, to sustain energy-consuming organs [20–22]. Conversely, BAT converts the products of endogenous lipolysis (or exogenous lipolysis from WAT) into heat, via a process called non-shivering thermogenesis [23,24]. Upon β-adrenergic receptor activation by NE in brown adipocytes, a protein called uncoupling protein 1 (UCP1) is activated by free fatty acids [23]. Active UCP1 provides a channel through which protons generated by oxidative phosphorylation can leak across the inner mitochondrial membrane. The resultant futile cycle of proton movement, and the engagement of other unrelated futile biochemical cycles, generates heat [25,26]. The third class of thermogenic UCP1-expressing adipocytes, known as beige [27] (or brite [28]) adipocytes also exists. These adipocytes are interspersed within WAT and exhibit intermediate phenotypes between white and brown adipocytes [7,27].

Non-shivering thermogenesis contributes to the maintenance of a stable core body temperature under ambient conditions that are lower than thermoneutrality (e.g. lower than 30°C in rodents). However, it also promotes energy balance in response to excessive caloric intake [23,29]. Notably, as the thermogenic activity of human BAT accounts for around 5 % of total metabolic rate, steady-state energy consumption by BAT has been proposed to account for weight loss in the order of 4 kg/year [30]. Consequently, much research has been dedicated to understanding how beige adipocytes could be recruited, or beige/brown adipocytes activated pharmacologically, to promote weight loss via elevated thermogenesis [31,32].

The metalloprotease ADAM17 [33,34] is an important shedding protease (‘sheddase’) that cleaves some prominent signaling molecules. ADAM17 substrates include cell adhesion molecules, the ectodomains of TNF and its cognate receptors, TNFRI & TNFRII, some ligands of the epidermal growth factor receptor (EGFR) and gp130, the IL-6 co-receptor whose cleavage by ADAM17 mediates IL-6-*trans* signaling [35,36]. Consequently, ADAM17 is the key protease in regulating inflammation and growth. It is implicated in pathologies associated with chronic inflammation [37], such as rheumatoid arthritis [38], and pathologies characterized by aberrant EGFR signaling, like some cancers [39].

While numerous studies have dissected the role(s) of ADAM17 in inflammation and growth control, very few studies have focused on the role of ADAM17 in energy balance and obesity [40,41]. However, although deletion of ADAM17 results in perinatal lethality, a fraction of mice survives until adulthood in some studies [40]. These KO mice were reported to exhibit hypermetabolism that was not compensated by their elevated food intake, and so exhibited a lean phenotype [40]. Similarly, in another study, mice with a heterozygous deletion of ADAM17 were protected from obesity (by elevating their energy expenditure in response to high-fat diet) and its associated metabolic dysregulation, such as glucose intolerance and insulin insensitivity [41]. This evidence suggests that ADAM17 plays an important role as a repressor of energy expenditure, via the proteolysis of as yet unidentified substrate(s). Although a subset of the phenotypes (e.g. improved glucose tolerance and insulin sensitivity) in ADAM17 mutant mice can be attributed to impaired TNF cleavage [42,43], the equal susceptibility of these TNF-null mice to obesity compared to WT mice [44] shows that the negative impact of ADAM17 on energy expenditure cannot be attributed exclusively to TNF[42,43]. Hence, the proteolytic substrate(s) via which ADAM17 negatively regulates energy expenditure remain to be identified.

Interestingly, the enhanced energy expenditure observed in ADAM17 mutant mice was associated with elevated levels of UCP1, implying that increased thermogenesis could underpin some of the observed hypermetabolic phenotypes [40]. In addition, we showed recently that mice null for inactive rhomboid 2 (iRhom2) are protected from diet-induced obesity [45]. Work from our group and others established that iRhom2 plays a crucial role in multiple aspects of ADAM17 biology including the control of its vesicular trafficking in the secretory pathway (which is coupled to the removal of its inhibitory prodomain) [46,47], its substrate specificity [48], and the activation of its activity on the cell surface by various stimuli [49,50]. In addition, an interactor called iTAP/FRMD8 is necessary to stabilize and maintain the function of the complex formed by iRhom2 and ADAM17 [51–53]. Notably, iRhom2-null mice also exhibit elevated energy expenditure and show increased thermogenesis (characterized by elevated BAT temperature)[45].

An early adaptation to an obesogenic dietary regime involves energy dissipation in the form of heat in BAT. This adaptation is driven by NE-mediated activation of β-adrenergic receptors. In the initial stage of adaptation, the sympathetic outflow to the adipose tissue is enhanced [54–56]. Therefore, the sympathetic nervous system (SNS) could impact ADAM17 activity and the shedding of its substrates. Indeed, agonists of G protein-coupled receptors have been shown to induce the activation of other signaling pathways, such as the EGFR, via the stimulation of ADAM17 activity [57]. This process is known as receptor transactivation. Binding of agonists such as angiotensin II, lysophosphatidic acid (LPA), ATP, serotonin and endothelin-1 to their cognate GPCRs activates cell surface proteases such as ADAM17 to shed their substrates, which in turn activate other signaling pathways [58,59].

Here, we hypothesized that the GPCR agonist NE could activate ADAM17 in adipocytes to shed substrate(s) that are important in the regulation of energy balance and specifically act to repress, thermogenesis. We show that deletion of ADAM17 in adipocytes is sufficient to elevate energy expenditure under steady state conditions and in obesity by increasing adipose tissue thermogenesis. This ameliorates overall organismal metabolic health in obese mice, by improving insulin sensitivity and glucose tolerance, dyslipidaemia, and hepatosteatosis. Moreover, proteomic studies in primary adipocytes revealed that in response to β-adrenergic receptor activation by NE, ADAM17 catalyzes the shedding of the cell surface signalling protein, Semaphorin 4B (SEMA4B). Our work identifies SEMA4B as a novel adipokine whose ADAM17-dependent release from adipocytes, in response to sympathetic outflow, acts as a negative regulator of thermogenesis by downregulating genes associated with lipid synthesis and catabolism.

## 2. Material and methods

### 2.1 Experimental animals

Mice were maintained in an SPF facility on a 12-hour light/dark cycle, at standard sub-thermoneutral conditions of 20-24 °C and an average of 50 % humidity, in ventilated cages with corn cob as bedding. All mice were housed in this condition, except stated otherwise. Mice had access to food and water *ad libitum*. Male mice were fed with normal chow or high-fat diet (SNIFF diet D12492 60 kJ% fat, E15742: 60% of energy from fat). Animals were routinely cohoused to homogenize differences in microbiota and other environmental conditions. In the event where mice were housed in different cages, bedding material was interchanged between cages to facilitate normalization of the microbiota.

To generate adipocyte specific *Adam17* KO mice (*Adam17*^adipoqΔ/Δ^), *Adam17*^fl/fl^ (loxP site flanking exon2 [60]) (Jackson Laboratory, 009567) were crossed with *Adipoq*^Cre^ mice [61]

All *in vivo* experiments were terminated by euthanizing mice with CO_2._ Tissue samples were thereafter collected, snap frozen in liquid nitrogen for storage at −80 °C for future processing or fixed in appropriate fixatives until they were ready to be processed.

### 2.2 Study approval

Animal procedures were approved by the national regulatory agency (DGAV – Direção Geral de Alimentação e Veterinária) and by the Ethics Committee of Instituto Gulbenkian de Ciência and the Institutional Animal Care (A012.2016 and A001.2020),and were carried out in accordance with the Portuguese (Decreto-Lei no. 113/2013) and European (directive 2010/63/EU) legislation related to housing, husbandry, and animal welfare.

### 2.3 Cold shock and experiment at thermoneutrality

Male mice were acclimatized in incubators set at the desired temperatures and appropriate humidity. For acute cold exposure, mice were moved from room temperature to incubator set at 4 °C for 6 hours. For acclimatization to cold (4 °C) or thermoneutrality (30 °C), mice were moved from 22 °C to the appropriate temperatures for at least 10 days.

### 2.4 Diet-induced obesity

To induce obesity, age-matched male mice were fed with HFD for 26 weeks. They were housed in large cages of 365 x 207 x 140 mm in dimension. These cages have the capacity to accommodate 16 mice. However, a maximum of 10 mice were housed per cage to guarantee enough space for the mice that develop obesity. Changes in mass of the mice were recorded weekly during the experiment.

### 2.5 Calorimetric measurements

The Promethion Core 8-cage system (Sable Systems) was used to assess body weight, food intake, energy expenditure (EE), locomotor activity (LA), and respiratory quotient (RQ) in mice fed on normal chow or HFD. Temperature was set at 22°C for all experiments except stated otherwise. For experiments in cold and thermoneutrality, temperatures were set as appropriate. Mice were acclimatised for at least 24 h in the metabolic cage system before the commencement of experiments.

### 2.6 Glucose and insulin tolerance tests

To test for glucose clearance or insulin sensitivity in the blood, mice exposed to normal chow or HFD for 22-24 weeks were fasted for 6 h (08:00 – 14:00). Fasting blood glucose was assessed by collecting blood from mice via tail puncture and using the one-touch Accu-Chek Aviva glucometer (Roche). Afterwards, mice were administered 1 g/Kg of glucose (Fisher) intraperitonially to test glucose clearance or insulin (Humulin) in a dosage of 0.8 UI/Kg to chow-fed animals or 1 UI/Kg to HFD-fed mice to test insulin sensitivity. Subsequent changes in blood glucose were measured at 15, 30, 60 90, and 120 min post glucose or insulin administration.

### 2.7 Enzyme-Linked Immunosorbent Assay (ELISA)

Blood was collected from 6 – 7 h fasted mice to quantify the concentration of insulin in the serum using an insulin ELISA kit (ALPCO). The medium of adipocytes was cleared and used to evaluate the levels of TNF, HB-EGF and Amphiregulin using commercial ELISA kits (88-7324-22 eBioscience, DY8239-05, R&D Systems, DY989, R&D Systems, respectively).

### 2.8 Serum and liver lipid profile

Serum from 6 - 7 h fasted mice were collected to quantify the levels of total cholesterol (TC), low density lipoprotein (LDLc), and triglycerides using enzymatic colorimetric kits (Spinreact). To quantify triglycerides (TG) in the liver, similar mass of liver was homogenised in a solution of chloroform and methanol (2:1). The homogenate was incubated overnight and centrifuged at 800 g for 15 min. The supernatant was subsequently mixed with 1/5 of its volume of saline solution (0.9% NaCl). The lower phase of the mixture was allowed to dry out after which a mixture of butanol/(Triton 100x/methanol) (2:1) was added for the colorimetric analysis of TG.

### 2.9 Histopathology

Mice tissues such as liver, adipose tissues, muscle, and pancreas were collected and weighed. Tissues to be processed for histology were fixed in 4% formaldehyde. Liver and muscle intended for oil red staining were fixed in 4 % formaldehyde for 24 h and thereafter transferred into 30% sucrose solution containing 0.005 % NaN_3_.

Adipose tissue (inguinal, epididymal, mesenteric, and retroperitoneal WAT, and BAT) and liver samples for H&E staining were embedded in paraffin, sectioned (3 μm per section) and stained with H&E. These sections were examined and captured using a Leica DMLB2 microscope and Leica DFC320 camera, respectively. Histopathological analysis was carried out by the in-house pathologist in a blinded manner.

H&E-stained adipocyte sections were converted into digital images and visualised in Nanozoomer-SQ and analysed in Fiji. Adipocyte size was determined using a macro on the H&E-stained adipocyte images.

In the liver, steatosis was scored from 0 – 4, representing <5 %, 5-33 %, 34-66 %, and >66 % of the total area of the section affected, respectively.

For oil red staining in the liver, samples previously stored in 30 % sucrose solution were snap-frozen and embedded in OCT. 8 μm of the tissues were cryosectioned and stained with oil red. Lipid content of the liver was scored as percentage of lipid per area of tissue.

### 2.10 Thermography

To measure BAT temperature, mice were anaesthetised using isoflurane. The fur covering the scapular and interscapular dermal surface was shaved. Mice were then allowed to recover by housing them individually in cages for 24 h with free access to food and water *ad libitum*. Food was withdrawn 2 h before thermal images of the interscapular region of the mice were captured using a thermal camera (FLIR). Resulting images were analysed in the FLIR image software. Rectal temperature was measured using a rodent thermometer (Bioseb).

### 2.11 Mouse primary adipocyte culture

The inguinal white adipose tissue (iWAT) and interscapular BAT (iBAT) were collected from 4–5-week-old male or female C57Bl/6, *Adam17*^fl/fl^, or *Adam17*^adipoqΔ/Δ^ mice for primary inguinal and brown adipocyte cultures, respectively. A culture was made from a pool of 3-5 mice. The tissues were cleaned, minced, and transferred into a 50 ml falcon. Two ml of Hank’s balanced salt solution (Biowest) supplemented with 1 mg/ml collagenase (Fisher Bioreagents), 10 mM CaCl_2_ (Sigma), and 2.2 mg/ml dispase II (Roche) was added to the minced tissue and put to shake at 37°C for 30-45 min. DMEMF12 (Biowest) was added to the mixture to stop digestion. The resultant solution was passed through 100 μm cell strainer. The filtrate was centrifuged at 500 g for 10 min at 4 °C. The pellet was resuspended again in DMEMF12 and filtered through 40 μm cell strainer. The resultant filtrate was again centrifuged at 500 g for 5 min at 4 °C. The resultant pellet was resuspended in complete medium (DMEMF12 supplemented with 1 % penicillin-streptomycin solution 100X, 2.5 μg/ml amphotericin B (Life Tech), 10 μg/ml gentamycin sulphate (Sigma), and 10% new-born calf serum (Biowest)), plated in 10 cm dish and incubated at 37°C, and 5% CO_2_. At confluence, preadipocytes were plated in collagen I (Corning) coated 6 or 12 well plates at 500,000, or 200,000 cells per well. respectively, to reach confluence. Differentiation was induced by exposing confluent cells to differentiation medium (complete medium supplemented with 1 mM 3-isobutyl methylxanthine (AppliChem), 2 μM rosiglitazone (Cayman), 10 μM dexamethasone (Cayman), 1 μg/ml insulin (Sigma), and 2 nM 3,3’,5-Triiodo-L-thyronine (Sigma)) on day 0 and day 2. On day 4 and day 6 post induction, differentiating cells were exposed to maintenance medium (complete medium supplemented with 1 μg/ml insulin (Sigma), and 2 nM 3,3’,5-Triiodo-L-thyronine (Sigma)). On day 8 post induction, the cells were used for relevant experiments.

### 2.12 Assessment of thermogenic gene expression in adipocyte cultures

Fully differentiated primary adipocytes were washed with serum free medium (SFM) (DMEMF12 supplemented with 1% penicillin-streptomycin solution 100X, 2.5 μg/ml amphotericin B (Life Tech), and 10 μg/ml gentamycin sulphate (Sigma)). SFM was replaced with SFM containing 2 % bovine serum albumin (BSA) (Merck). To stimulate the βARs, the cells were treated with 2 μM norepinephrine (NE) (Sigma). The cells and/or medium were collected at specific time points depending on the experiment.

Adipocyte stimulation with TNF was performed by pre-treating differentiated WT adipocytes with vehicle, versus recombinant TNF (10 ug/ml, 410-MT-010/CF, R&D) for 2h. After this, cells were treated with norepinephrine (NE, at 0.4 uM) for further 6 hours and RNA was isolated.

For the experiment showing increased shedding of ADAM17 substrates upon βAR stimulation, fully differentiated WT and ADAM17 KO primary adipocytes are treated with NE for 1 h. For experiments aimed at assessing the regulation of thermogenic genes in WT and ADAM17 KO primary adipocytes fully differentiated cells were treated with NE for 6 h, while cells intended for assessing protein expression were treated with NE for 3 h, 6 h, 12 h, and 24 h, depending on the specificities of the experiment.

### 2.13 Quantitative transcriptional analysis

Adipose tissues (iBAT, iWAT, and epididymal WAT), liver, and primary adipocytes collected at the end of experiments for transcriptional analysis were snap frozen and lysed in NR buffer of the NZYTech total RNA extraction kit. Subsequent steps for collection and purification of RNA were guided by the protocol of the kit. First-strand cDNA was synthesised from RNA using the Xpert cDNA synthesis master mix (GRiSP). SYBR green or Taqman reagents (GRiSP) were used for real time PCR. The comparative C_T_ method was employed and GAPDH and Actin were used in normalising gene expression. The primer sequences for qPCR and Taqman probes used are listed below.

### 2.14 Evaluation of expression of Sema4B and its receptors in human tissues

The data in Figures 5I and J were replotted from publicly available data. The data for Figure 5I were replotted from data from an experiment in which two human pluripotent stem cell lines (H9 and KOLF2-C1) were differentiated into brown adipocytes [62]. The data in Figure 5J were obtained from a study in which human adipocytes were differentiated from SVF obtained from subcutaneous WAT and subclavicular BAT from human subjects [63].

**Figure 5:**
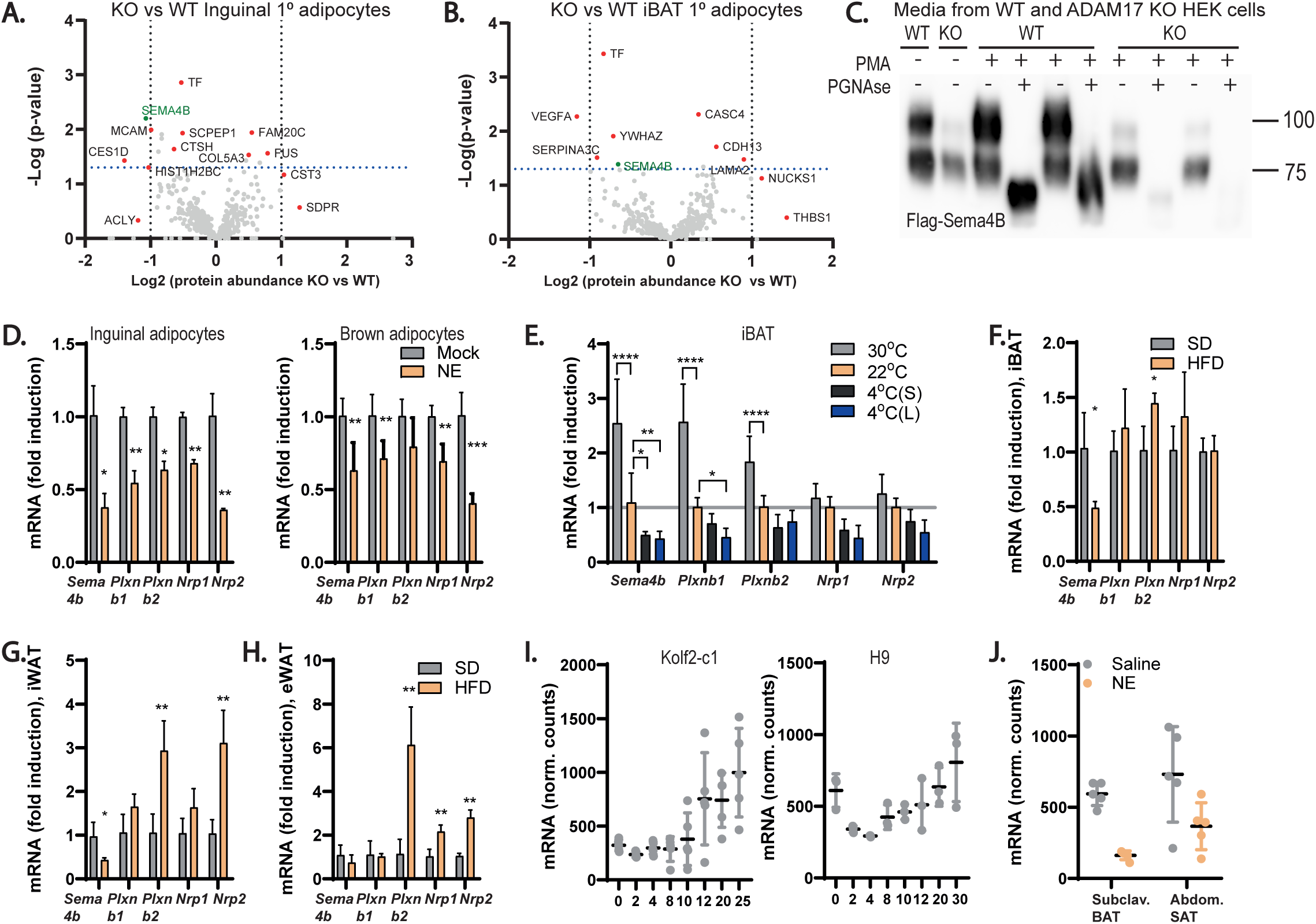
Semaphorin 4B is a novel adipokine and an ADAM17 substrate. **(A-B)**, Volcano plot depicting proteins detected in the secretome of ADAM17 KO vs WT inguinal (**A**) or brown (**B**) primary adipocytes. Proteins with log2 difference greater than or less than zero are detected at higher or lower levels, respectively in conditioned medium from ADAM17 KO adipocytes. Proteins above the horizontal cut-off (-Log pvalue=1.3) are significantly detected at levels higher or lower levels in conditioned medium from ADAM17KO adipocytes (n=5). **(C)**, Immunoblot of Sema4B in conditioned media from WT and ADAM17 KO Hek293 cells transfected with full-length Sema4B plasmid. Cells were either untreated or treated with 1μM of phorbol myristate acetate (PMA) and deglycosylated using PNGAse (n=4). **(D-E)**, mRNA levels of *Sema4b* and its receptors *Plxnb 1 & 2, Nrp 1 & 2* in inguinal and brown primary adipocytes treated without or with 2μM of NE (**D**) (n=5), and in the interscapular BAT of mice exposed to different ambient temperatures; thermoneutrality (30 ^°^C), room temperature (22 ^°^C), acute cold exposure (4^°^ S) (4 ^°^C for 6 hours), and chronic cold exposure (4^°^ L) (4 ^°^C for 10 days) (**E**) (n=6). **(F-H)**, mRNA levels of *Sema4b* and its receptors; *Plxnb* 1 & 2, *Nrp* 1 & 2 in the interscapular BAT (**F**), inguinal WAT (**G**), and epididymal WAT (**H**) of lean and HFD-induced obese mice (n=4). **(I)**, Expression (counts) of *Sema4B* mRNA over the course of the differentiation of human-derived stem cells, Kolf2-c1 and H9 to matured adipocytes. **(J)**, Expression (counts) of *Sema4b* mRNA in human subclavicular BAT and abdominal subcutaneous WAT stimulated without or with 2μM of NE. Results presented as mean ± SD. *P<0.05, **P<0.01, ***P<0.001, ****P<0.0001.

### 2.15 Western Blotting

Mouse adipose tissues and primary adipocytes samples were lysed in lysis buffer (1% Triton X-100, 150 mM NaCl, 50 mM Tris-HCl, pH 7.4) supplemented with protease inhibitors and 10 mM 1,10 phenanthroline. To improve the detection of ADAM17, homogenates were incubated overnight in concanavalin A resin at 4°C. The incubated homogenates were then centrifuged at 1000 g for 2 min at 4°C. The pellets (resin) were washed in lysis buffer (supplemented with protease inhibitors and 10 mM 1,10 phenanthroline) 3 times. The beads were eluted in sample buffer supplemented with 15% sucrose by heating at 65°C for 15 min. When homogenates were not enriched to improve the detection of specific proteins, equal amount of protein from different samples were denatured in sample buffer by heating at 65 °C for 15 min. In some cases, the denatured lysates were digested for 2 hr at 37 °C with the deglycosylating enzyme PNGase F, which removes high mannose N-linked glycans added in the ER and complex N-linked glycans found in the later secretory pathway. Then, the samples were denatured for 15 min at 65°C. The samples were fractionated in SDS-PAGE before being transferred onto PVDF membrane. The membranes were blocked in Tris-buffered saline with 0.1 % Tween® 20 Detergent (TBS-T) supplemented with 5 % milk or BSA for at least 30 min and then incubated in primary antibodies (diluted in TBS-T supplemented with 5 % milk or BSA) (1:1000) overnight at 4°C. The following primary antibodies were used: rabbit anti-ADAM17 antibody (1:1000, Ab39162, Abcam), mouse anti-Flag HRP (1:1000; A8592; Sigma), mouse anti-Transferrin receptor (1:1000, 13-6800, Life Technologies). The membranes were subsequently washed 4 times in TBS-T for 10 min before incubating in the appropriate secondary antibody anti-rabbit HRP (1:5000; 1677074P2, Cell Signaling Technology) or anti-mouse HRP (1:5000; 1677076P2, Cell Signaling Technology) (diluted in TBS-T supplemented with 5 % milk or BSA) (1:5000) for at least 30 min. The membranes were again washed 4 times in TBS-T for 10 min followed by a 10 min wash in Phosphate buffer saline (PBS). Protein bands were detected using ECL. Images were captured using the Amersham^TM^ imager 600.

### 2.16 Primary adipocyte secretome analysis with high-performance secretome protein enrichment with click sugars (hiSPECS)

To analyse the secretome from *Adam17*^fl/fl^ and Adam17*^Adipoq^/^^* primary adipocytes, maintenance medium was added to differentiated adipocytes on day 6 post-induction of differentiation (300,000 preadipocytes were plated per well of a 6-well plate before the induction of differentiation) was supplemented with 50 μM of ManAZ (Thermo Fisher Scientific, Cat #C33366) for 48h. The conditioned medium from these cells was collected to analyse the secretome of the primary adipocytes in steady state. However, to assess the changes in the secretome upon induction of thermogenesis, the 48 h-long maintenance medium supplemented with 50 μM of ManNAz was replaced with SFM supplemented with 10 % serum and 50 μM of ManNAz for 12 h. The cells were stimulated, or not, with 2 μM of norepinephrine during this period. 1 mL of the medium from the cells were collected and filtered through a 0.45 μm centrifuge tube filter (Costar® Spin-X #10649351). Proteinase inhibitor (1:500) was added to the filtrate and the samples were stored at −20 °C until further use. The hiSPECS workflow, as previous described in detail [64] was performed. In brief, glycoproteins were enriched using Concanavalin A and azido-modifed proteins were immobilized on magnetic DBCO-beads (Jena Bioscience #CLK-1037-1) via click chemistry. Stringent washing steps reduced contaminants and peptides were released from the beads by tryptic digest. Desalted peptides were analysed using liquid chromatography mass spectrometry (LC-MS/MS) using an EASY-nLC 1200 UHPLC system (Thermo Fisher Scientific) coupled to an Q Exactive HF mass spectrometer (Thermo Fisher Scientific). Data-dependent acquisition (DDA) and data-independent acquisition (DIA) analysis was applied and data analysis was performed with MaxQuant and Spectronaut. Proteins were considered sufficiently expressed if detectable in at least 4 of the 5 samples per treatment condition. To enrich for candidate membrane proteins shed by metalloprotease activity we focused on proteins that have a single transmembrane helix or a glycosylphosphatidylinositol (GPI) anchor, as previously described[65]. A p-value ≤ 0.05 was used as cut-off for significantly and differentially shed proteins.

### 2.17 Assessment of Sema4B cleavage in cultured cells

Wild type and ADAM17 KO HEK293ET cells were kind donations from the Becker-Pauly Lab, University of Kiel, Germany. Full length flag-tagged Sema4B plasmid was donated by the Püschel Lab, University of Munster, Germany [66]. To test for the potential shedding of Sema4B ectodomain by ADAM17, Wild type and ADAM17 KO HEK293 cells were plated overnight in a 6-well plate (500,000 cells per well). The day after, the cells were transfected with Sema4B or an empty vector using Polyethyleneimine (PEI). In brief, cells plated overnight had their medium changed for 2 ml of complete medium containing 2.5 μg of Sema4B plasmid DNA or empty vector and 7.5 μg of PEI per well. Twenty-four hours after transfection, the medium of the cells was aspirated, the cells were washed with SFM, and serum-starved for 4 h. Thereafter, cells were treated with vehicle or phorbol-12-myristate-13-acetate (PMA) for 1 h to stimulate ADAM17 activity. The media were collected to precipitate protein content and the cells were lysed to be processed for western blot as previously described[67].

### 2.18 Lentiviral transduction

HEK293 ET packaging cells (6 x 10^6^) were transfected with pCL-Eco packaging plasmid[68] plus SV40 Large T Ag, or (16.6 x 10^6^) with the pMD-VSVG envelope plasmid, psPAX2 helper plasmid plus pLEX, or pLEX containing mouse truncated Semaphorin 4B cDNA fused to a Flag tag. The packaging vectors for the production of lentivirus were described previously [69]. WT brown preadipocytes from a WT newborn mouse was immortalized as previously described[70]. In summary, isolated cells were split in two and seeded in a 35-mm plate at 40% confluence and one well was transduced with 2ml of SV-40 supernatant supplemented with polybrene 8 μg/mL twice with a 24h interval. The non-transduced cells were used as control. Then, the immortalized cells were transduced with 20 μl of 150x ultracentrifuged (90,000 g, 4 h, at 4 °C) empty vector (pLex), Semaphorin 4B viral supernatant (resuspended in 0.1% BSA in PBS) supplemented with polybrene 8 μg/mL, and selected with puromycin (10 μg/mL) for 2 days to generate stable cell lines. These cells were differentiated using the standard protocol.

### 2.19 Expression and purification of Semaphorin 4B ectodomain

Expi293F cells (Thermo Fischer Scientific) were cultured in Expi293 Expression Medium and transfected with the pCMVi-SV40 ori-based plasmid encoding amino acids 1 to 700 of mouse Semaphorin 4B (Uniprot ID Q62179) followed by PreScission cleavage site, hexahistidine tag and Twin Strep tag. Five days after transfection, medium with secreted ectodomain of Sema4B was collected, and flash frozen in liquid nitrogen. For purification of Sema4B ectodomain, the thawed medium was loaded onto Strep-Tactin® Superflow® high capacity cartridge (IBA Lifesciences GmbH) pre-equilibrated in the binding buffer (100 mM Tris-HCl pH 8.0, 150 mM NaCl). Unbound material was washed away with 10 column volumes of binding buffer, and Sema4B ectodomain was eluted with 5 column volumes of the elution buffer (100 mM Tris-HCl pH 8.0, 150 mM NaCl, 2.5 mM desthiobiotin). Glycerol was added to the purified protein to 10% (v/v) before flash freezing it in liquid nitrogen. Finally, purified Sema4B ectodomain was depleted from any possible traces of endotoxins by the Pierce™ High Capacity Endotoxin Removal Spin Column (Thermo Fischer Scientific), according to the manufacturer’s protocol. The final preparation of Sema4B ectodomain was flash frozen in small aliquots in liquid nitrogen and stored at −80°C for further use.

### 2.20 RNA sequencing and data analysis

Immortalized murine brown adipocytes stably tranduced with lentiviruses harboring pLEX empty vector or a mimetic of the ADAM17-cleaved form of Sema4B (described above, 2.18) were plated at a density of 300,000 cells/well in 6-well plates. The following day, after achieving confluence, the cells were differentiated to matured adipocytes over a period of 8 days as described above (2.11). Medium from matured adipocytes was aspirated and the cells were washed with serum-free medium (SFM). Afterwards, the cells were incubated in SFM supplemented with 2% BSA with or without 2 μM NE for 6 h. At the end of this timepoint, the cells were washed with cold PBS and the cells were lysed using the NR buffer of the NZYTech total RNA extraction kit. Subsequent steps for collection and purification of RNA were guided by the protocol of the kit. For the RNA seqeuncing, full-length cDNAs were generated following the SMART-Seq2 protocol described by Picelli *et al*., 2014 [71]. After quality control using Fragment Analyzer (Agilent Technologies), library preparation including cDNA fragmentation, PCR-mediated adaptor addition and amplification of the adapted libraries was done following the Nextera library preparation protocol (Nextera XT DNA Library Preparation kit, Illumina), as previously described [72]. Libraries were confirmed by Fragment Analyzer (Agilent Technologies) and then sequenced (NextSeq2000, Illumina) using 100 SE P2. Sequence information was extracted in FastQ format, using Illumina DRAGEN FASTQ Generation v3.8.4. Library preparation and sequencing were optimized and performed by Genomics Unit at the Instituto Gulbenkian Ciência. Raw fastq reads were processed using FastQ Screen to identify any non-murine reads prior to alignment. Reads passing were aligned to the mouse genome (mm10, GRCm38.p4) using STAR v2.7.9a [73]. Counts per gene were calculated using htseq-count[74]. Differential gene expression analysis was performed with DESeq 2[75], with an adjusted *p-*value cut-off of ≤0.05 and a foldchange cut-off of ≥ 1. Pathway enrichment was performed with DAVID [76]. Normalised gene counts were log2-scaled and z-scored for data visualisation. Heatmaps were generated in the R statistical programming language (https://cran.r-project.org/).

## 3. Results

### 3.1 Beta-adrenergic receptor activation stimulates ADAM17 to repress thermogenesis in adipocytes

Our previous study established that mice null for iRhom2, the key regulator of ADAM17 trafficking and activity [46,47] exhibit elevated adipose tissue thermogenesis and Ucp1 levels [45]. As noted above, a fraction of ADAM17 KO mice that escape embryonic lethality and survive into adulthood also exhibit elevated Ucp1 levels [40]. Together, this evidence led us to hypothesize that ADAM17 acts within adipose tissue to repress thermogenic signals. To test this hypothesis, we generated mice in which ADAM17 was deleted in adipose tissue using adiponectin-cre (adipoq-cre) [61] (**Fig. S1A**). ADAM17 activity can be activated by certain classes of G protein-coupled receptors (GPCR)[77,78]. As adipocyte beta adrenergic receptors (βARs) control thermogenic responses and lipolysis [79], we tested the hypothesis that the endogenous βAR agonist norepinephrine (NE) can transactivate pathways mediated by substrates of ADAM17. Hence, we isolated adipocyte progenitors from the stromal vascular fraction (SVF) from inguinal adipose tissues of control floxed (ADAM17^fl/fl^) versus *Adam17*^adipoqΔ/Δ^ mice in which ADAM17 was deleted in adipose tissues (henceforth referred to as “WT” and “KO” throughout) and differentiated them to mature adipocytes *ex vivo*. As shown in **Fig. S1A**, a significant decrease was observed in the protein levels of ADAM17 in adipocytes differentiated from SVF obtained from the KO mice relative to the WTs. Focusing on iWAT, a tissue that is susceptible to beiging [27], mature primary adipocytes were stimulated with NE and the shedding of several crucial ADAM17 substrates (TNF (Tumor necrosis factor); HB-EGF (Heparin-binding EGF); and AREG (Amphiregulin)) into the culture media was quantified by ELISA. Notably, NE stimulation significantly increased the shedding of TNF, HBEGF, and AREG in WT adipocytes, while shedding of these substrates was inhibited in the ADAM17 KO adipocytes (**Fig. 1A**). These data show that activation of endogenous βARs triggers a cascade of events that stimulate the proteolysis of ADAM17 substrates and links ADAM17 regulation to a critical pathway that governs adipocyte physiology.

**Figure 1:**
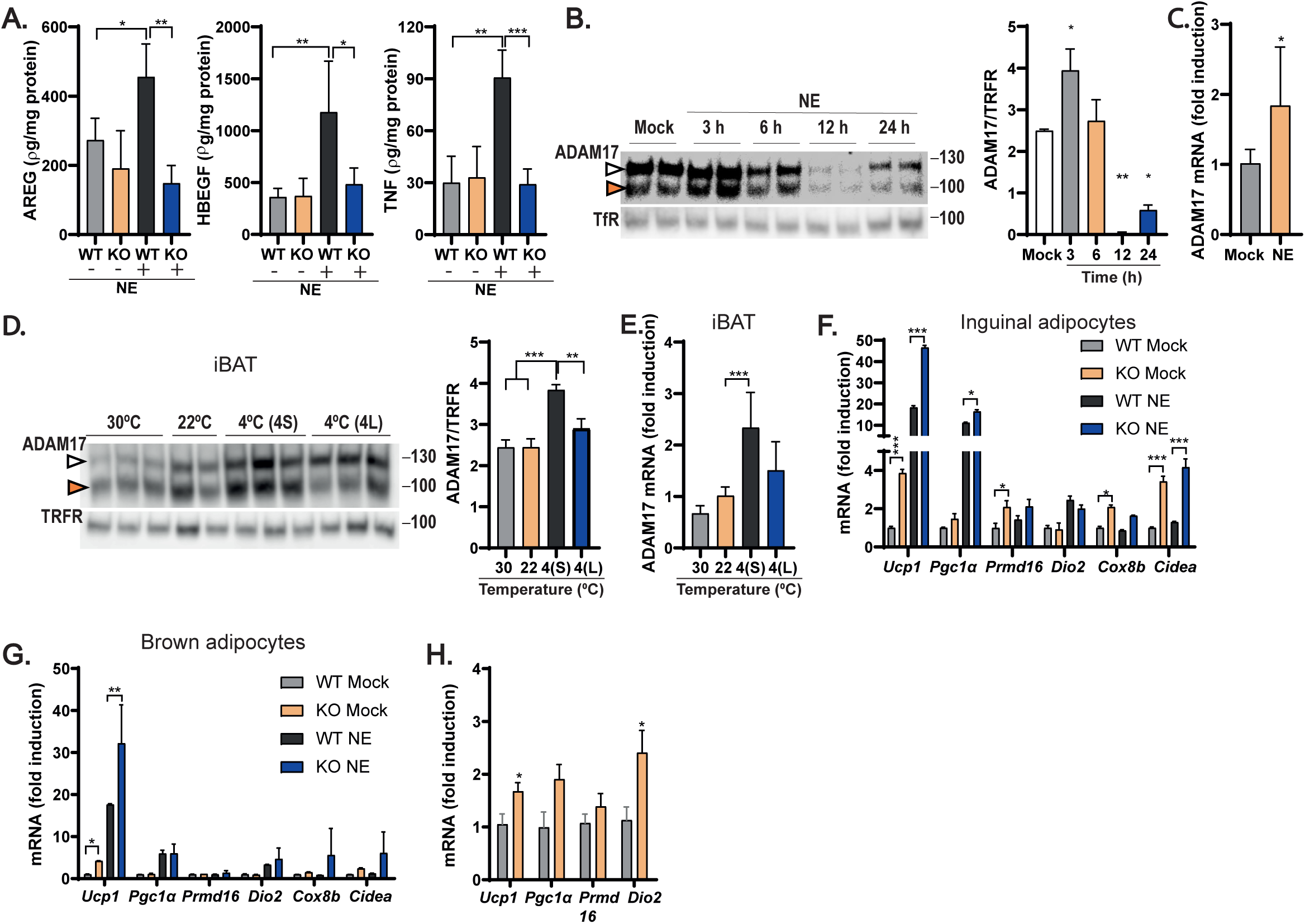
Loss of ADAM17 in adipocytes increases thermogenesis in primary adipocytes and elevates energy expenditure *in vivo*. **(A)**, Amphiregulin (AREG), heparin-binding growth factor (HB-EGF), and tumor necrosis factor (TNF) levels quantified by ELISA in conditioned media from control and ADAM17 KO primary adipocytes with or without stimulation with 2μM of norepinephrine (NE) (n=4). **(B)**, Immunoblots showing mature (100 KDa) (orange arrowhead) and immature (130 KDa) (white arrowhead) ADAM17 from untreated WT primary adipocytes and primary adipocytes stimulated with 2μM of NE over different time points for a total period of 24 h. Total ADAM17 (matured and immature ADAM17) quantified as a ratio of the loading control, transferrin receptor (TfR) (n=4). **(C)**, mRNA level of *Adam17* in WT primary adipocytes without or with treatment with 2μM of NE (n=4). **(D)**, Immunoblot showing mature (100 KDa) (white arrowhead) and immature (130 KDa) (orange arrowhead) ADAM17 levels in interscapular brown adipose tissue (iBAT) from WT mice exposed to different ambient temperatures; thermoneutrality (30 ^°^C), room temperature (22 ^°^C), acute cold exposure (4^°^ S) (4 ^°^C for 6 hours), and chronic cold exposure (4^°^ L) (4 ^°^C for 10 days). Total ADAM17 (matured and immature ADAM17) quantified as a ratio of the loading control, transferrin receptor (TfR) (n=6 all conditions except n=4 for 22 ^°^ C). **(E)**, mRNA level of *Adam17* in the iBAT of mice from D (n=6). **(F-G)**, mRNA level of positive regulators of thermogenesis in control and ADAM17 KO primary inguinal (**F**) and brown (**G**) adipocytes treated without or with 2μM of NE (n=4). **(H)**, mRNA level of the thermogenic genes, *Ucp1, Pgc1α, Prmd16*, and *Dio2* in the iBAT of lean control (WT) and adipocyte-specific ADAM17 KO (KO) mice (n=5). Results presented as mean ± SD. *P<0.05, **P<0.01, ***P<0.001, ****P<0.0001.

In addition to the stimulatory effects of NE in promoting the shedding of some ADAM17 substrates, we also observed that βAR activation also impacted the levels of ADAM17. As shown in **Fig. 1B**, stimulation of WT mature primary inguinal adipocytes with NE over a 24-hour period revealed a complex impact of βAR activation on ADAM17. At 3 hours post-stimulation, βAR activation triggered an increase in the overall levels of ADAM17, particularly in the levels of the mature form of the protease that has reduced molecular weight (**Fig. 1B**). At later timepoints, the levels of ADAM17 decreased over time becoming almost non-detectable 12 hours post-stimulation with NE, after which, a gradual recovery in ADAM17 levels was observed 24 hours post induction with NE (**Fig. 1B**). βAR activation also elevated *Adam17* mRNA levels by approximately 2-fold in primary inguinal WT adipocytes (**Fig. 1C**). To test the impact of these observations in an *in vivo* setting, we modulated sympathetic neuron output and hence NE release and βAR activation by exposing WT mice to different ambient temperatures. Mice were either acclimated to thermoneutrality (30 °C, to minimize sympathetic output), room temperature (22 °C), or cold (4 °C, to enhance sympathetic output) after which interscapular brown adipose tissues were isolated. As shown in **Figure 1D-E**, exposure of WT mice to acute cold increased the protein and mRNA expression levels of ADAM17 in brown adipose tissue. Taken together with the primary adipocyte data (**Fig. 1A-C**), this suggests that, in response to sympathetic output and βAR activation, ADAM17 could play a role in the physiological adaptation of the BAT to decreasing ambient temperature.

To understand the impact of ADAM17 on thermogenesis, we quantified the level of expression of thermogenic genes in primary inguinal and brown adipocytes differentiated from the SVF of WT versus KO mice. Notably, loss of ADAM17 in adipocytes upregulated the expression of some thermogenic genes in inguinal and brown adipocytes, both under basal conditions and in response to NE (**Fig. 1F-G**). Consistent with this, *in vivo,* ADAM17 KOs maintained on a standard diet expressed higher levels of thermogenic genes such as *Ucp1*, *Pgc1α*, *Prmd16*, and *Dio2* in their interscapular BAT than their WT counterparts (**Fig. 1H**). Our data indicate that *Adam17* mRNA and protein levels and sheddase activity are modulated by the neurotransmitter NE to repress thermogenesis. This led us to investigate the hypothesis that ADAM17 could serve as a rheostat, licensed by adrenoceptor stimulation, to limit the overactivation of the thermogenic programs in adipocytes.

### 3.2 ADAM17 deletion in adipocytes is anti-obesogenic and promotes increased energy expenditure via elevated non-shivering thermogenesis

The observation of elevated expression of thermogenic genes in primary adipocytes and in the interscapular BAT from lean mice, together with the previous observation that ADAM17 KO[40] or iRhom2-deficient mice [45] have a hypermetabolic phenotype could imply that loss of ADAM17 drives enhanced energy expenditure via non-shivering thermogenesis *in vivo*. To test this, we assessed metabolic parameters such as body weight, food intake, and energy expenditure in our WT and KO mice maintained on a standard diet. As shown in **Fig. S1B-C**, no significant differences were observed in the body mass and food intake of WT and KO mice. Notably however, the KO mice showed a significant increase in total energy expenditure and oxygen (O_2_) consumption and carbon dioxide (CO_2_) production during the dark phase (**Fig. S1D-F**) which are consistent with the reported hypermetabolic phenotype observed in whole-body ADAM17 KO mice [40]. Like O_2_ consumption and CO_2_ production, respiratory exchange ratio (RER), a measure of the relative utilization of carbohydrates versus fats as a fuel source [80] is elevated only in the dark phase in the KO mice (**Fig. S1G**). This enhanced energy expenditure in the KO mice is not accounted for by differences in locomotion between WT and KO mice (**Fig. S1H**).

With the observation of elevated energy expenditure in the KO mice especially in the dark phase (**Fig. S1D**), we predicted that the KO mice should be protected from obesity since these animals should be better able to expend excess calories. To test this, we allowed WT and KO mice to feed *ad libitum* on a high-fat diet (HFD) for 26 weeks (**Fig. S2A**). Notably, the KO mice gained less body weight and accumulated less fat in adipose tissues relative to the WT mice (**Fig. 2A-B**; **Fig. S2A-D**), despite consuming more calories than their WT counterparts (**Fig. 2C**). This increase in food intake was previously observed in lean ADAM17 global KO mice[40]. To assess the overall metabolic phenotypes of these mice and to account for the reduced weight gain in the KOs, we placed the obese mice in metabolic cages and quantified energy expenditure, oxygen consumption, CO_2_ production, and overall locomotor activity. To control for the differences in body weight between WT and KO mice on HFD in our study of energy expenditure, oxygen consumption and CO_2_ production, we analyzed our data using analysis of covariance (ANCOVA), introducing body weight as a covariate[81] As shown in **Fig. 2D-F**, the KO mice expended more energy, consumed more oxygen, and produced more CO_2_ relative to the WT mice suggesting that protection of KO mice from obesity is driven by increased metabolic rate. While the respiratory quotient (**Fig. 2G**) and locomotor activity (**Fig. 2H**) of KO mice were unremarkable, notably, we observed a significant increase in BAT thermogenesis (**Fig. 2I**). As the core body temperature of KOs was unaltered compared to WT mice, (**Fig. 2J**), we conclude that under an obesogenic regime, KO mice are hypermetabolic and expend significantly more energy on thermogenesis than their WT counterparts. These data support the hypothesis that ADAM17 is an endogenous inhibitor of thermogenesis.

**Figure 2:**
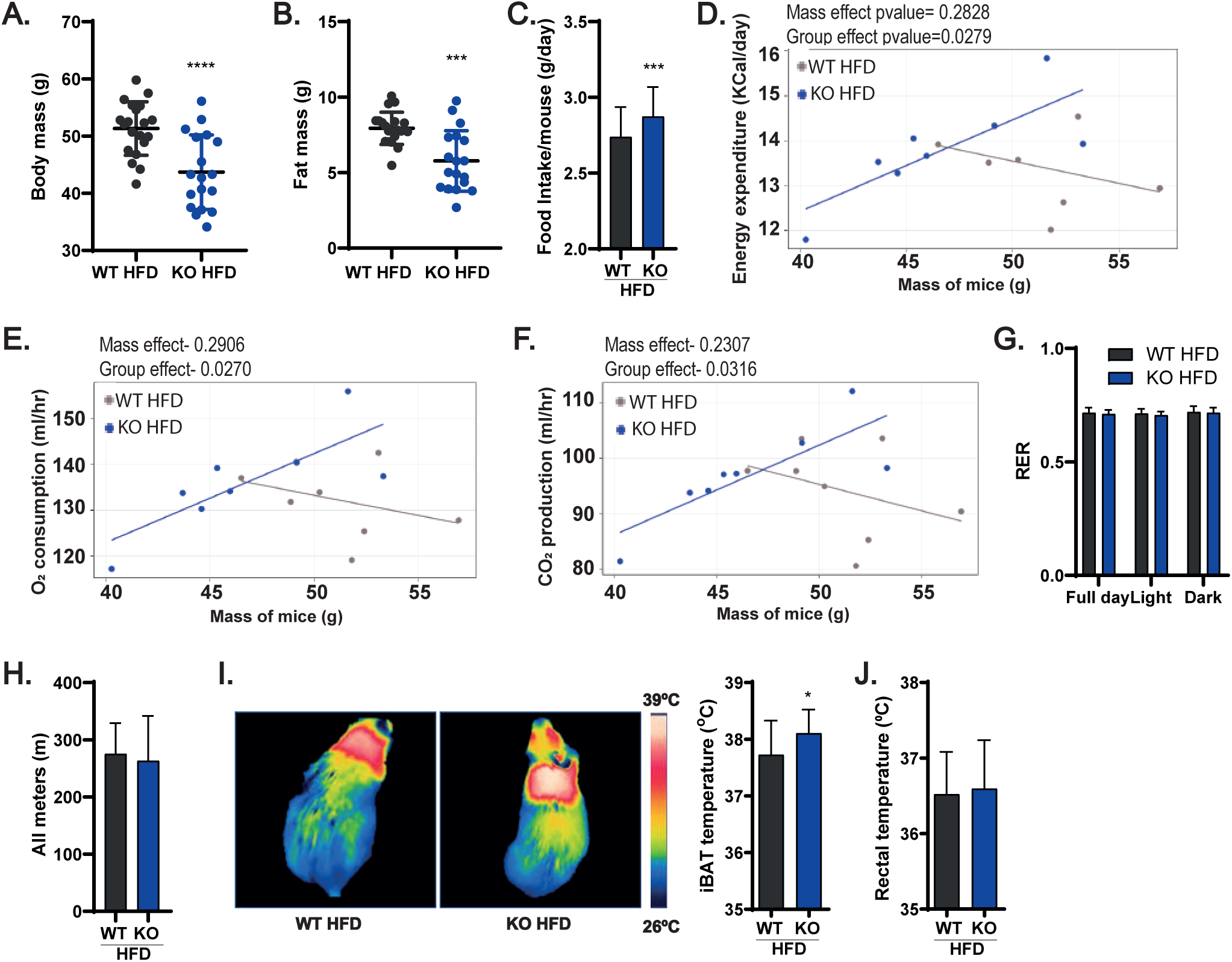
Mice null for ADAM17 in adipocytes are less prone to developing diet induced obesity by increasing energy expenditure. **(A-C)**, Body mass (**A**), pooled fat mass (visceral adipose tissues, inguinal adipose tissue, and iBAT) (**B**), and mean daily food intake per mouse (**C**) of WT and KO mice on high-fat diet (HFD) for 26 weeks (WT n=20, KO n=18). **(D-F)**, Scatter plots of energy expenditure (**D**), oxygen consumption (**E**), and carbon dioxide production (**F**) over 24 h in obese WT and KO mice with body mass used as a covariate (WT n=8, KO n=7). **(G-H)**, Respiratory exchange ratio (RER) (**G**), and distance covered (**H**) over 24 h period of WT and KO mice on HFD for 26 weeks (WT n=8, KO n=7). (**I-J**), Thermographic representation and quantification of temperature of the interscapular region (**I**), and rectal temperature (**J**) of WT and KO mice after 26 weeks on HFD (WT n=16, KO n=17). Results presented as mean ± SD. *P<0.05, **P<0.01, ***P<0.001, ****P<0.0001

### 3.3 ADAM17 deletion protects from obesity-associated adipocyte hypertrophy in some fat depots by enhancing the expression of genes involved in lipid catabolism and thermogenesis

As KO mice were protected from obesity and exhibited an overall reduction in fat mass, we next sought to establish which adipose depots were most affected. We found that except for the epididymal WAT (**Fig. 3A**) the visceral adipose tissues (mesenteric and retroperitonial WATs) (**Fig. 3B, C**), the inguinal WAT (**Fig. 3D**), and the interscapular BAT (**Fig. 3E**) were significantly reduced in mass in the KOs compared to WT controls. Moreover, for some fat depots tested, we found that the KO mice exhibited reduced adipocyte area (**Fig. 3G, I-J**). Consistent with our previous data showing elevated thermogenic gene expression in ADAM17-deficient BAT (**Fig. 1H**), upon exposure to HFD, the BAT of KO mice better retained its characteristic multilocular lipid droplet morphology with increased LD number per unit area (**Fig. 3K**) compared to WT controls.

**Figure 3:**
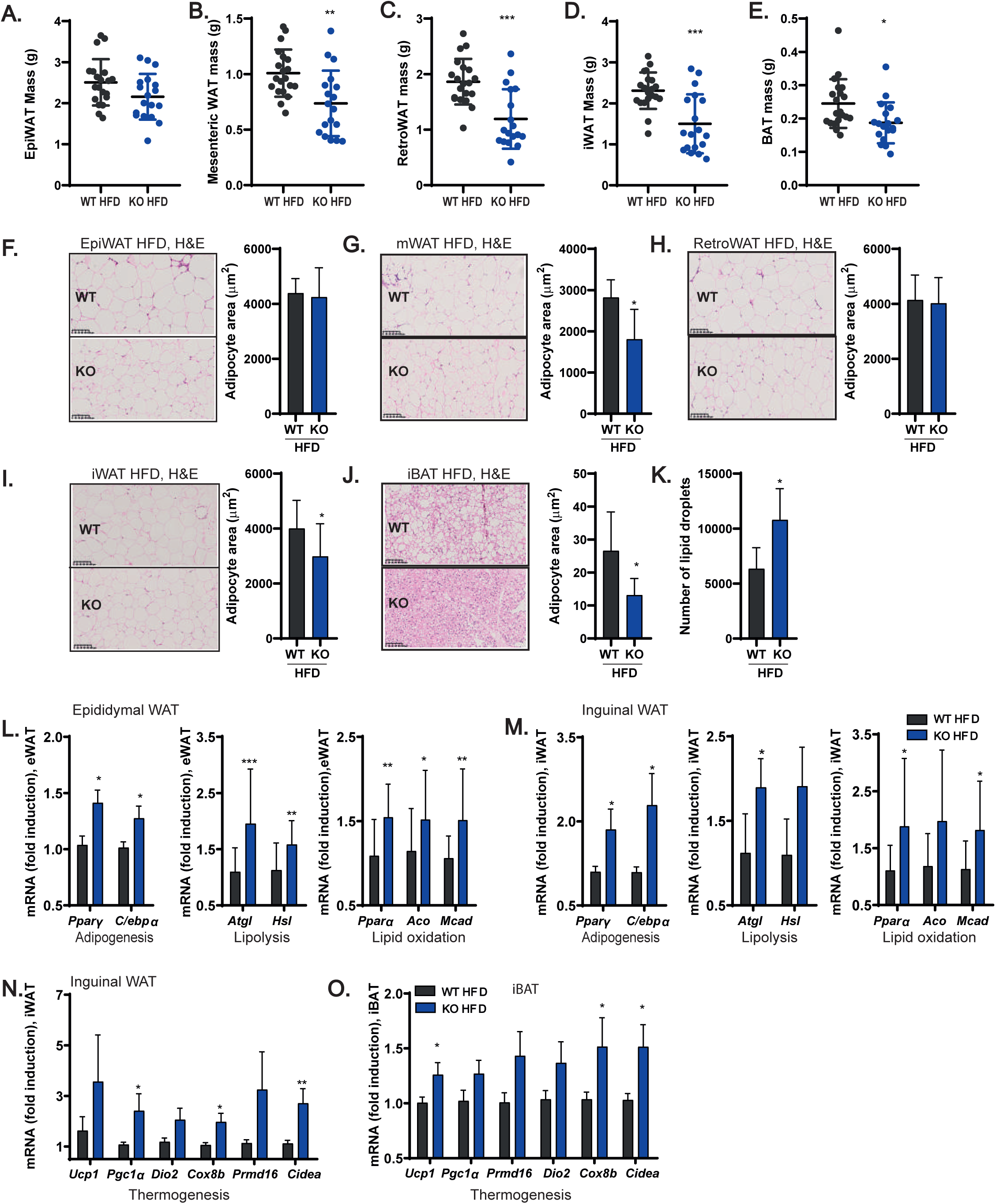
Mice null for ADAM17 in adipocytes are protected from white adipocyte hypertrophy and brown adipocyte whitening on HFD. **(A-E)**, Mass of epididymal (**A**), mesenteric (**B**), retroperitoneal (**C**), inguinal (**D**), and interscapular (**E**) brown adipose tissues of WT and KO mice after 26 weeks of HFD (WT n=20, KO n=18). **(F-J)**, H&E staining and quantification of adipocyte size (area) from epididymal (**F**) (WT n=11, KO n=13), mesenteric (**G**) (WT n=6, KO n=6), retroperitoneal (**H**) (WT n=7, KO n=7), inguinal (**I**) (WT n=12, KO n=12), and interscapular brown (**J**) (WT n=6, KO n=6) adipose tissues of WT and KO mice on HFD. **(K)**, Quantification of lipid droplets from H&E stained iBAT slides of WT and KO on HFD (WT n=6, KO n=6). **(L)**, mRNA levels of adipogenic (WT n=12, KO n=12), lipolytic (WT n=20, KO n=18), and lipid oxidation (WT n=19, KO n=17) related genes in the epididymal adipose tissue of WT and KO mice on HFD. **(M)**, mRNA levels of adipogenic (WT n=18, KO n=15), lipolytic (WT n=18, KO n=15), and lipid oxidation (WT n=18, KO n=15) related genes in the inguinal adipose tissue of WT and KO mice on HFD. **(N-O)**, mRNA level of thermogenic genes in the inguinal (**N**) (WT n=17, KO n=15) and interscapular brown (**O**) (WT n=18, KO n=14) adipose tissue of WT and KO mice on HFD. Results presented as mean ± SD. *P<0.05, **P<0.01, ***P<0.001, ****P<0.0001

We further analyzed how ADAM17-deficient white adipose tissues (epididymal and inguinal) better adapt to an obesogenic diet than their WT counterparts. Adipose tissue expansion is due to two mechanisms: hyperplasia (an increase in adipocyte numbers within the tissue) versus hypertrophy (an increase in adipocyte cell size)[82]. Notably, the KO WATs expressed significantly higher levels of mRNA of the adipogenic genes *Pparγ* and *C/ebpα* that promote adipose tissue hyperplasia, relative to the WT controls (**Fig. 3L-M**). In addition, there was an increase in the mRNA levels of the lipolytic genes *Atgl* and *Hsl* in the WATs from the KO mice (**Fig. 3L-M**). An increase in adipogenesis (hyperplasia) and elevated lipolysis could account for the decreased adipose tissue mass encountered in the KO mice.

The ratio of the lipolysis products released into circulation versus those oxidized and liberated as heat within adipose tissues is critical in determining whether lipids are ectopically deposited in other metabolic organs, such as the liver[83]. Notably, ADAM17-deficient WATs have elevated expression of genes involved in lipid oxidation (**Fig. 3L-M**) while elevated expression of thermogenic genes was found in the inguinal WAT and interscapular BAT of the KO mice (**Fig. 3N-O**). Hence, the potentially elevated partitioning of ingested lipids towards catabolism, and BAT thermogenesis and inguinal WAT beiging could spare the KO mice from adiposity and ectopic fat accumulation.

### 3.4 Loss of ADAM17 protects from obesity-driven hepatic lipid spillover and insulin resistance

Consistent with the notion of better lipid handling implied by the elevated expression of genes associated with lipid oxidation, the KO mice on HFD were significantly protected from obesity-induced liver enlargement (**Fig. 4A**) and lipid accumulation (**Fig. 4B-C**), associated with reduced liver triglyceride accumulation compared to WT controls (**Fig. 4D**). To further understand whether the protection from hepatosteatosis in KOs results from a reduced propensity for this organ to uptake, store, or export lipids, we quantified the mRNA expression of genes involved in these processes. As shown in **Fig. S3A**, the expression of the fatty acid transporter *Cd36* was significantly downregulated in the livers of KO mice compared to WT controls, possibly because of their leaner phenotype, as the expression of CD36 is promoted during obesity[84; 85]. Furthermore, consistent with the improved metabolic health of the KO animals, genes involved in the positive regulation of lipolysis and lipid oxidation (**Fig. S3B-C**) were upregulated in the liver of KO mice compared to WT controls. In addition, we found an upregulation of the mRNA levels of transcripts that regulate the formation and export of lipoproteins from the liver, such as *Apob1* and *Apoe,* in KO livers compared to WT controls (**Fig. S3D**). Together, these results show that loss of ADAM17 in adipose tissues protects from lipotoxicity-derived non-alcoholic fatty liver disease (NAFLD) development typified by increased hepatic lipid uptake, and decreased oxidation and export. [84].

**Figure 4:**
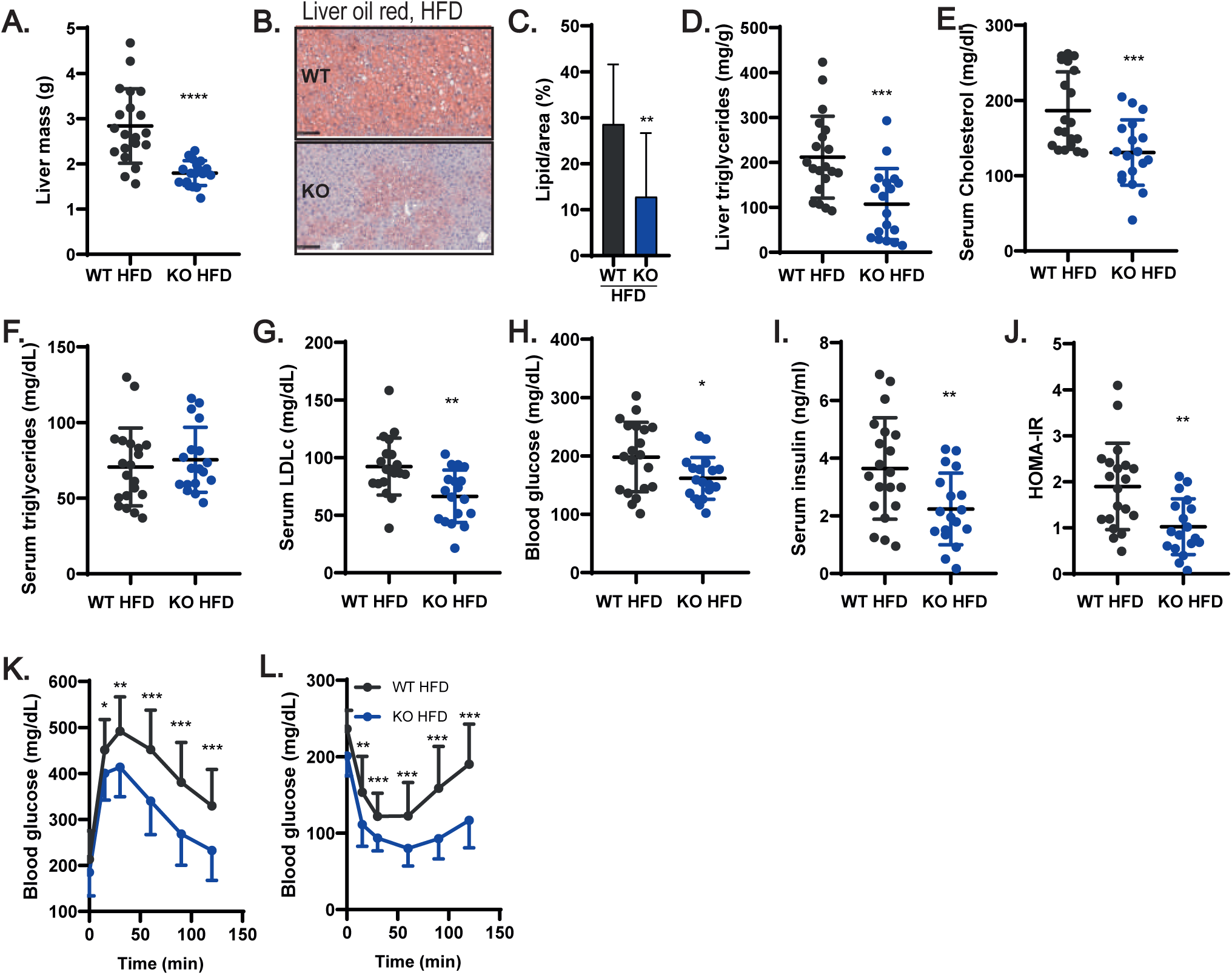
Mice null for ADAM17 in adipocytes are protected against HFD induced ectopic lipid deposition in the liver, dyslipidaemia, and systemic insulin resistance. **(A-C)**, Mass of liver (**A**) (WT n=20, KO n=18) and images of oil-red stained liver samples (**B**) (WT n=13, KO n=13), and quantification of percentage of lipid-laden area of oil-red stained liver samples (**C**) (WT n=13, KO n=13), from WT and KO mice after 26 weeks of HFD. **(D)**, Triglyceride content of liver from WT and KO mice on HFD (WT n=20, KO n=18). **(E-G)**, Fasting serum levels of total cholesterol (**E**), triglycerides (**F**), and low-density lipoprotein-cholesterol (LDLc) (**G**) in WT and KO mice on HFD (WT n=20, KO n=18). **(H-I)**, Fasting blood glucose (**H**) and serum insulin levels (**I**) in WT and KO mice on HFD (WT n=20, KO n=18). **(J-L)**, Homeostatic assessment model for insulin resistance (HOMA-IR) (**J**), glucose (**K**) and insulin (**L**) tolerance tests in WT and KO mice on HFD (WT n=20, KO n=18). Results presented as mean ± SD. *P<0.05, **P<0.01, ***P<0.001, ****P<0.0001

Consistent with the notion that the capacity of the liver to handle lipids has an important impact on the serum lipid profile and upon metabolic health in general, we observed significantly lower levels of total cholesterol and LDL-cholesterol in the serum of KO mice compared to WT animals fed on HFD (**Fig. 4E&G**), with no change in serum triglyceride levels (**Fig. 4F**). Because of lower levels of circulating cholesterol associated with the observed healthier adipose tissues (**Fig 3A-M**), KO mice exposed to obesogenic conditions presented lower fasting glycaemia and insulinaemia, and were more insulin sensitive (**Fig. 4H-J**). The subjection of these mice to glucose or insulin tolerance tests confirmed that the KO mice on HFD are more glucose tolerant and insulin sensitive, respectively, relative to obese WT mice (**Fig. 4K-L**), indicating that the KO mice were protected from obesity-associated glucose metabolism alterations. Notably, no differences were observed in the lipid profile (**Fig. S3E-G**), fasting glycaemia and insulinaemia (**Fig. S3H-I**) and insulin and glucose tolerance (**Fig. S3J-K**) in KO *vs* WT mice maintained on standard diet (SD). Hence, loss of ADAM17 in adipose tissues has the most noticeable impact within the context of positive energy balance associated with an obesogenic diet.

### 3.5 Semaphorin 4B is a novel ADAM17 substrate expressed in adipocytes and regulated by thermogenic stimuli and obesity

Our results so far show that ADAM17 ablation in adipocytes protects from the metabolic alterations associated with an obesogenic diet by promoting energy expenditure in adipose tissues. This predicts that proteolytic shedding of substrate(s) by ADAM17 from the cell surface of adipocytes should drive signaling pathways that negatively regulate energy expenditure. As stated earlier, the major pathways regulated by ADAM17 are the TNFR and the EGFR signaling pathways. However, neither of these pathways has been compellingly shown to contribute to adipose tissue thermogenesis or weight gain upon exposure to HFD[42–44; 86; 87] and their reported roles in metabolic diseases are often contrasting[88]. Another critical pathway mediated by ADAM17 proteolysis, IL-6 *trans*-signaling, exerts its salutary metabolic effects centrally by reducing food intake and by promoting energy expenditure [89]. The action of IL-6 hence promotes weight loss and decreases body fat [89–92]. Although our adipose tissue-specific ADAM17 mutants exhibited elevated food intake(**Fig. 2C**) they are nonetheless hypermetabolic and less susceptible to weight gain on HFD, suggesting that defective IL-6 *trans*-signaling does not underpin the basis of the anti-obesogenic and hypermetabolic phenotype observed in our model.

We therefore adopted an objective proteomic approach to interrogate the secretome from WT versus ADAM17 KO primary inguinal and brown adipocytes to identify ADAM17 substrates secreted in thermogenic adipocytes. We used hiSPECs (high-performance secretome protein enrichment with click sugars), an approach that enables the specific labelling and enrichment of glycoproteins by culturing cells with an azido group-containing sugar [93]. This approach carries the advantage that cells can be grown under normal serum-containing conditions [93]. As shown in **Figure 5A-B**, after filtering for potentially shed proteins[65] (see Materials and Methods), we observed a significant reduction in the levels of the cleaved form of a transmembrane protein called Semaphorin 4B (Sema4B) in the secretome from the KO inguinal and brown primary adipocytes (**Fig 5A-B**). Semaphorins are secreted, transmembrane and cell-surface-attached proteins that signal mainly by interaction with Plexin receptors, and act on axonal guidance, and regulate morphology and motility of many cell types [94]. We confirmed that Sema4B is a genuine novel ADAM17 substrate that can be shed in response to PMA, by comparing the supernatant from WT versus ADAM17 KO HEK cells transfected with full length Sema4B (**Fig. 5C**).

A key feature of genes involved in thermal regulation (e.g. *Ucp1*) is the modulation of their transcriptional levels by thermogenic cues in response to sympathetic outflow and consequent β-adrenoceptor activation [95]. Intriguingly, we found that the mRNA levels of *Sema4b* and its known receptors (Plexins) and co-receptors (Neuropilins [96]) were significantly downregulated in response to NE stimulation of primary inguinal and brown mouse adipocytes (**Fig. 5D**). Furthermore, the levels of *Sema4b* and its associated receptors/co-receptors correlated with exposure of mice to ambient temperature, decreasing in expression levels from thermoneutrality to 4 °C (**Fig. 5E**), which is potentially consistent with a negative role in thermogenesis regulation. In obese mice, the expression of *Sema4B* was also downregulated in interscapular BAT and inguinal WAT but not in epididymal WAT (**Fig. 5F-H**). However, the expression of its receptors and coreceptors was upregulated in these fat depots in obese animals (**Fig. 5F-H**). Furthermore, the analysis of datasets on human stem cells differentiated into mature brown adipocytes[62] showed that the expression of Sema4B is regulated during adipogenesis (**Fig. 5I**), and that the treatment of adipocytes differentiated from human subclavicular BAT or abdominal subcutaneous WAT with NE[63] downregulates the expression of *Sema4b* (**Fig. 5J**), as we observed in primary mouse adipocytes (**Fig. 5D**). Moreover, gene set enrichment analysis (GSEA) from the datasets from Din *et al*.[97] showed that enhanced BAT thermogenesis (e.g. increased *Ucp1* expression) in response to ingestion of mixed carbohydrate diet negatively correlates with genes involved in semaphorin and plexin signaling (**Fig. S4A**). Taken together, the murine and human data indicate that *Sema4b* expression levels are significantly affected by ambient temperature, diet-induced thermogenesis, or NE, suggesting that SEMA4B is implicated in the regulation of thermogenesis.

### 3.6 Sema4B is a negative regulator of thermogenesis

As ADAM17 cleaves Sema4B and thermogenic cues in multiple contexts can modulate *Sema4B* mRNA levels, we next tested the effects of cleaved Sema4B upon thermogenic gene expression in a range of models. Strikingly, we observed that stable expression of a cleaved mimetic of Sema4B in immortalized WT brown adipocytes significantly inhibited *Ucp1* expression in naïve and NE-stimulated mouse adipocytes (**Fig. 6A**). Brown adipocyte differentiation is important for determining the thermogenic capacity of adipocytes while lipolysis is essential for Ucp1 activation. Therefore, these two key processes may be impacted by Sema4B to repress thermogenesis. Indeed, Sema4B repressed the expression of the major lipolytic genes, *Atgl/Pnpla2* and *Hsl* (**Fig. 6B**). Sema4B also repressed expression of the adipogenic genes, *Ppar*γ and *C/ebpα* (**Fig. 6C**). Furthermore, in independent experiments, mammalian-produced recombinant soluble Sema4B ectodomain similarly repressed *Ucp1* levels in primary brown adipocytes in response to NE (**Fig. 6D**), downregulated the expression of lipolytic genes (**Fig. 6E**), but had a minimal effect on adipogenic ones (**Fig. 6F**). Hence, our data suggest that ADAM17, through Sema4B cleavage, secretion and autocrine action on adipocytes, downregulates key processes required to support thermogenesis.

**Figure 6:**
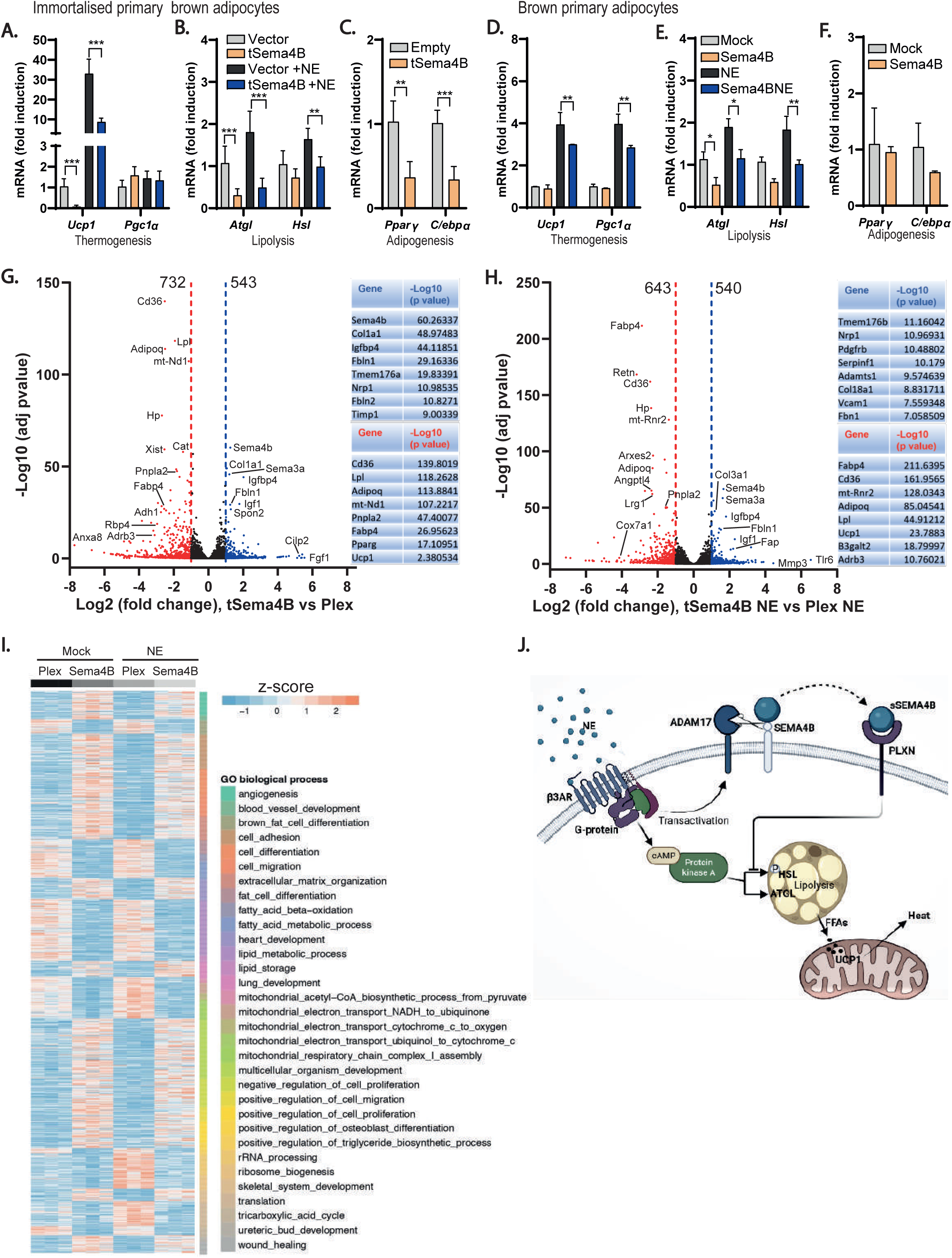
Semaphorin 4B regulates thermogenesis by repressing UCP1 expression. **(A-C)**, mRNA expression of the thermogenic, lipolytic, and adipogenic genes in immortalised primary adipocytes transduced with empty vector or truncated Sema4B and treated with or without NE. (n=6). **(D-F)**, mRNA expression of the thermogenic, lipolytic, and adipogenic genes in primary adipocytes treated with or without recombinant Sema4B and subsequently treated with or without NE (n=5). **(G)**, Volcano plots showing differentially expressed genes in immortalized primary brown adipocytes transduced with truncated sema4B (tSema4B) or empty vector (Plex) and a list of some upregulated and downregulated genes by tSema4B (n=3). **(H),** Volcano plots showing differentially expressed genes in immortalised primary brown adipocytes transduced with truncated sema4B (tSema4B) or empty vector (Plex) in response to NE and a list of some upregulated and downregulated genes by tSema4B in response to NE (n=3). **(I),** Heat map of upregulated and downregulated genes by tSema4B in naïve and NE-stimulated cells according to the GO:term, biological process. **(J)**, Schematic representation of the potential mechanism via which Sema4B represses thermogenesis in adipocytes. Results presented as mean ± SD. *P<0.05, **P<0.01, ***P<0.001, ****P<0.0001.

To better understand the impact of Sema4B on adipocyte biology in general and specifically to elucidate how Sema4B perturbed processes required to support thermogenesis, we performed RNAseq on WT immortalised primary adipocytes overexpressing cleaved Sema4B or the empty vector, under unstimulated (**Fig. 6G**) versus NE-stimulated conditions (**Fig. 6H**). Under unstimulated conditions, when a threshold of a log2 fold change of at least 1 or -1 was applied, 543 and 732 genes were upregulated and downregulated (**Fig. 6G**). Notably, the cohort of genes that were upregulated by Sema4B included Nrp1 (a semaphorin receptor that has been reported to bind to Sema4B) plus genes associated with remodeling of the extracellular matrix (Col1a; Fbln1, Fbln2, Timp1). By contrast, prominently downregulated hits were genes involved in lipid metabolism and fatty acid update such Lpl (lipoprotein lipase) the fatty acid transporter CD36 and the fatty acid binding protein Fabp4 and lipolysis (*Atgl/Pnpla2*). As anticipated from the qPCR studies, we also observed downregulation of genes associated with BAT thermogenesis (*Pparγ, Ucp1*). In NE-treated cells, Sema4B expression upregulated and downregulated 540 and 643 genes, respectively (with a log2 fold change of at least 1 or −1) (**Fig. 6H**). Similar to unstimulated samples, soluble Sema4B-overexpressing NE-treated cells upregulated hits included mRNAs involved in extracellular matrix remodeling (*Col18a1, Serpinf1, Adamts1,Vcam1*) and vasculogenesis (*Fbn1* and *Nrp1*). As for mock-treated cells, downregulated hits included genes associated with fatty acid uptake (*Fabp4, Cd36, Lpl*), lipolysis *(Pnpla2/Atgl)* and BAT thermogenesis (*Ucp1, Adrb3*) (**Fig. 6H**). When gene enrichment analysis was applied on all differentially expressed genes that passed the p ≤ 0.05 threshold, amongst the genes repressed by Sema4B under unstimulated conditions, the biological processes represented included brown adipocyte differentiation; fatty acid import, storage and beta oxidation (**Fig. 6I**). When NE-stimulated, the downregulated hits additionally included genes associated with mitochondrial respiration, while the upregulated hits were enriched in mRNAs associated with extracellular matrix organization, development, and angiogenesis. Our transcriptomic data emphasize that cleaved Sema4B represses several key processes that are required to support thermogenesis.

## 4. Discussion

One of the most interesting findings to emerge from the present study is the connection between β-adrenoceptors, a cornerstone of adipocyte physiology, and ADAM17. NE treatment of primary adipocytes stimulates shedding of ADAM17 substrates, modulates ADAM17 expression, while exposure of mice to decreasing ambient temperature (which triggers NE release) upregulates ADAM17 levels. Notably, the effects on ADAM17 behaviour observed in our model are similar to the response to the ADAM17 stimulant, PMA, where an initial increase in ADAM17 activity and maturation is followed by a subsequent decrease in expression due to its degradation [98].

Our data identify a negative regulatory loop (**Fig. 6J**) whereby “licensing” of thermogenesis by β-adrenoceptor activation triggers an autocrine pathway in adipocytes, driven by ADAM17-mediated shedding of Sema4B, a novel adipokine. Soluble Sema4B triggers a transcriptional response in adipocytes that represses the expression of several genes that control processes that are required to support thermogenesis (e.g. lipid uptake, lipolysis) and *Ucp1* expression itself. Speculatively, when triggered by β-adrenoceptor activation, this novel mechanism, which has similarities to EGFR transactivation [99], may fine-tune thermogenic responses or act as a negative feedback loop to prevent untrammelled resource depletion to avoid a sustained state of negative energy balance. Speculatively, this pathway may represent a metabolic vulnerability within the context of obesity. Future studies will be required to determine whether the Sema4B pathway is actionable for the improvement of metabolic health.

Our study reveals that ablation of ADAM17 in adipocytes promotes the expression of thermogenic genes (and elevates thermogenesis) in the iBAT of animals on HFD and on SD under standard housing conditions, leading to improved metabolic health of animals on an obesogenic diet. This is reminiscent of the hypermetabolic phenotype reported for the fraction of whole-body ADAM17 KOs that escape perinatal lethality [40]. Our adipose tissue-specific ADAM17-deficient mice also had a marginal increase in energy expenditure on SD, although this appears not to confer easily observable metabolic benefits (e.g. upon glucose homeostasis and lipid profile) relative to WT mice under the sub-thermoneutral conditions of standard animal housing [100–102].

The increased energy expenditure exhibited by ADAM17 KOs could be accounted for by several inter-related mechanisms that enhance the adipose tissue beiging and thermogenesis observed in the KO mice (**Fig. 1F-H; Fig. 2I-J; Fig. 3N-O**) and consequently, an enhanced catabolic state consistent with elevated levels of mRNAs associated with lipolysis or beta-oxidation (**Fig. 3L-M**). Our cellular **(Fig. 1, Fig. 6)** and transcriptomic studies **(Fig. 6)** support the view that this is an adipocyte-autonomous circuit that represses lipid uptake and catabolism and thermogenesis (**Fig. 6J**). However, it will be interesting to determine, in future studies, whether adipocyte ADAM17 can also impinge on sympathetic tone, the sympathetic innervation of adipose tissues, the recruitment/function of adipose tissue-associated immune cells, or indeed upon endocrine cross talk between adipose tissues and other organs (e.g. the brain, liver).

Although there is no prior established connection between Sema4B and metabolic regulation, we can make several general inferences concerning how Sema4B could negatively regulate thermogenesis from the biology of other semaphorins. Semaphorins can signal through intracellular kinases such as PKA [103], which are required for β-adrenergic signal transduction (e.g. to regulate lipolysis). However, the functions of semaphorins are not linear and seem to be determined by the tissue microenvironment. Adding to this complexity is the ability of semaphorins to bind not only to their multiple cognate receptors plexins and neuropilins, but also to several other receptors and coreceptors (e.g. CD72, Tim2, integrins, and proteoglycans[94; 104–107]). Indeed, several receptor tyrosine kinases (e.g. VEGFR2, ErbB2, and Met) associate with plexins and neuropilins and are transactivated upon semaphorin binding [94]. Aside from acting as ligands, semaphorins can also act as receptors, via a phenomenon known as reverse signaling [94].

Consistent with our earlier observations of elevated thermogenesis and hypermetabolism in ADAM17 KOs (e.g. a context where Sema4B cleavage/secretion is frustrated), our transcriptomic data suggest that cleaved Sema4B could impair thermogenesis in a pleiotropic manner. For example, this includes impinging directly upon thermogenic gene expression, or upon metabolic processes that are required to support thermogenesis (e.g. lipolysis, lipid uptake). Sema4B also negatively impacts on beta-adrenoceptor expression, which would also indirectly dampen thermogenesis. Interestingly, Sema4B also reduced the mRNA levels of the transcription factors C/ebpα and Pparγ which play crucial roles in (brown) adipocyte differentiation[108,109]. Aside from repressing genes involved in lipolysis, Sema4B seems to negatively impact all steps involved in lipid uptake until their breakdown into energy substrates to fuel heat generation by brown adipocytes.

Semaphorins act, in part, by regulating the activation state of GTPases [94]. Guanine nucleotide exchange proteins (GEFs) keep GTPases turned on by promoting their binding to GTP while GTPase-activating proteins (GAPs) promote the binding of GTPase to GDP [110]. Interestingly, plexin receptors have a conserved GAP homology domain that activates the GTPase activity of the Rap and Ras family of GTPases [111]. Speculatively, in the context of adipocytes, plexin activation by Sema4b could increase the GAP activity of the G-proteins associated with β-ARs, maintaining the G-proteins in an inactive state. This would dampen downstream signal transduction in response to NE, lipolysis and UCP1 activation. Future studies are needed to address this interesting possibility.

In addition to the pronounced negative impact on expression of mRNAs associated with lipid homeostasis and thermogenesis, cleaved Sema4B also promoted the upregulation of extracellular matrix (ECM)-associated genes including transforming growth factor *beta* (Tgf-β), a fibrosis-promoting cytokine[112] and various collagens. Indeed, ADAM17 has an established pro-fibrotic activity [113–115] while pro-fibrotic roles for semaphorins have recently emerged in a variety of disease contexts including in the lung, kidney and the cornea [116–119]. Interestingly, several studies have implicated the aberrant expression of extracellular matrix-associated genes, including collagens, with defective adipose tissue remodeling during obesity [120]. However, evidence for semaphorin-mediated promotion of adipose tissue fibrosis is limited to a single study on Sema3C [121], while the impact of pro-fibrotic genes in adipocyte thermogenesis is still poorly understood [122,123]. Speculatively, cleaved Sema4B could act on pre-adipocytes or mature adipocytes to promote fibrosis, to trigger a range of negative impacts on adipocyte physiology like on adipose tissue plasticity and insulin resistance [121] or thermogenesis [124].

Our current knowledge on the role of semaphorins in metabolism is very limited and centered on the Sema3 class of semaphorins. To date, the most prominent example that implicates a semaphorin in obesity is on the role of class 3 semaphorins, of which rare variants have been isolated from severely obese human subjects [125]. These semaphorins act upon neuropilin-2 in the hypothalamus to promote the development of melanocortin neuronal circuits that regulate energy homeostasis; loss of neuropilin-2 in pro-opiomelanocortin neurons reduces energy expenditure and promotes weight gain [125]. Moreover, Sema3A is expressed in adipocytes together with neuropilin-1 and is negatively regulated during cold acclimation in BAT [126,127]. In addition, Sema3G has opposing roles in adipogenesis: its plasma levels are increased in obese patients, while its deletion protects mice from HFD-induced weight and fat gain, insulin resistance and glucose tolerance and liver lipogenesis [128,129]. Although our present study identifies a novel adipocyte-autonomous axis for semaphorin signaling in metabolic regulation, it will be interesting to determine whether soluble forms of Sema4B can act hormonally on energy centers in the brain (or indeed, whether shedding of other semaphorins by ADAMs plays a role in energy regulation in the brain).

Although the metalloproteases involved in the reported proteolysis of other semaphorins remain to be fully delineated, interestingly, the ectodomains of class 3-7 semaphorins [130–132] or their receptors [133,134] can be cleaved by ADAMs or other metalloproteases. Sema3C for example was shown to be cleaved by ADAMTS1 [135], while Sema3A signaling is regulated by the proteolysis of neuropilin-1 by ADAM17 and ADAM10 [133]. We have shown here that Sema4B is cleaved by ADAM17. However, this does not rule out the involvement of other proteases in Sema4B proteolysis (in other tissues), nor indeed, novel metabolic regulatory roles for other semaphorins potentially cleaved by ADAM17 or other ADAMs.

ADAM17 is a pleiotropic enzyme crucial for the development and progression of numerous diseases. As targeting ADAM17 has been demonstrated to cause toxicity because of its pleiotropic activity and because of the cross-reactivity of ADAM17 inhibitors with similar metalloproteases [136,137], finding new targets for specific branches of ADAM17 biology is an imperative for the development of novel ADAM17 associated therapies. Therefore, as a novel ADAM17-shed adipokine, Semaphorin 4B could be a promising candidate for the development of new therapeutic strategies to treat obesity.

## 5. Conclusions

Our work establishes an important role for ADAM17 as a negative regulator of adipocyte thermogenesis that is regulated by beta-adrenoceptor signalling. It also identifies Sema4B as a novel ADAM17-shed adipokine that acts in an adipocyte-autonomous manner to limit uncontrolled energy consumption during thermogenesis.

## Acknowledgements

The authors thank the Ethics Committee, Animal Facility, Histopathology and Flow cytometry units and the Antibody Service of the Instituto Gulbenkian de Ciência. We thank Hana Dvořáková and Zuzana Vaitová for assistance with production of recombinant Semaphorin 4B. We thank Andreas Püschel for the Sema4b expression plasmid. We acknowledge the support of Fundação Calouste Gulbenkian; Queen’s University Belfast; Worldwide Cancer Research (14–1289); a Marie Curie Career Integration Grant (project no. 618769); Fundação para a Ciência e Tecnologica (FCT) grants, SFRH/BCC/52507/2014, PTDC/BEX-BCM/3015/2014 and LISBOA-01–0145-FEDER-031330), funding from ‘La Caixa’ Foundation under the agreement <LCF/PR/HR17/52150018>; support from the ERDF/ESF project ChemBioDrug (No. CZ.02.1.01/0.0/0.0/16_019/0000729) and by the Deutsche Forschungsgemeinschaft (DFG, German Research Foundation) under Germany’s Excellence Strategy within the framework of the Munich Cluster for Systems Neurology (EXC 2145 SyNergy–ID 390857198). This work was developed with the support of the research infrastructure Congento, project LISBOA-01–0145-FEDER-022170, co-financed by Lisboa Regional Operational Programme (Lisboa 2020), under the Portugal 2020 Partnership Agreement, through the European Regional Development Fund (ERDF), and Foundation for Science and Technology (Portugal).

---

**Supplementary Figure 1:**
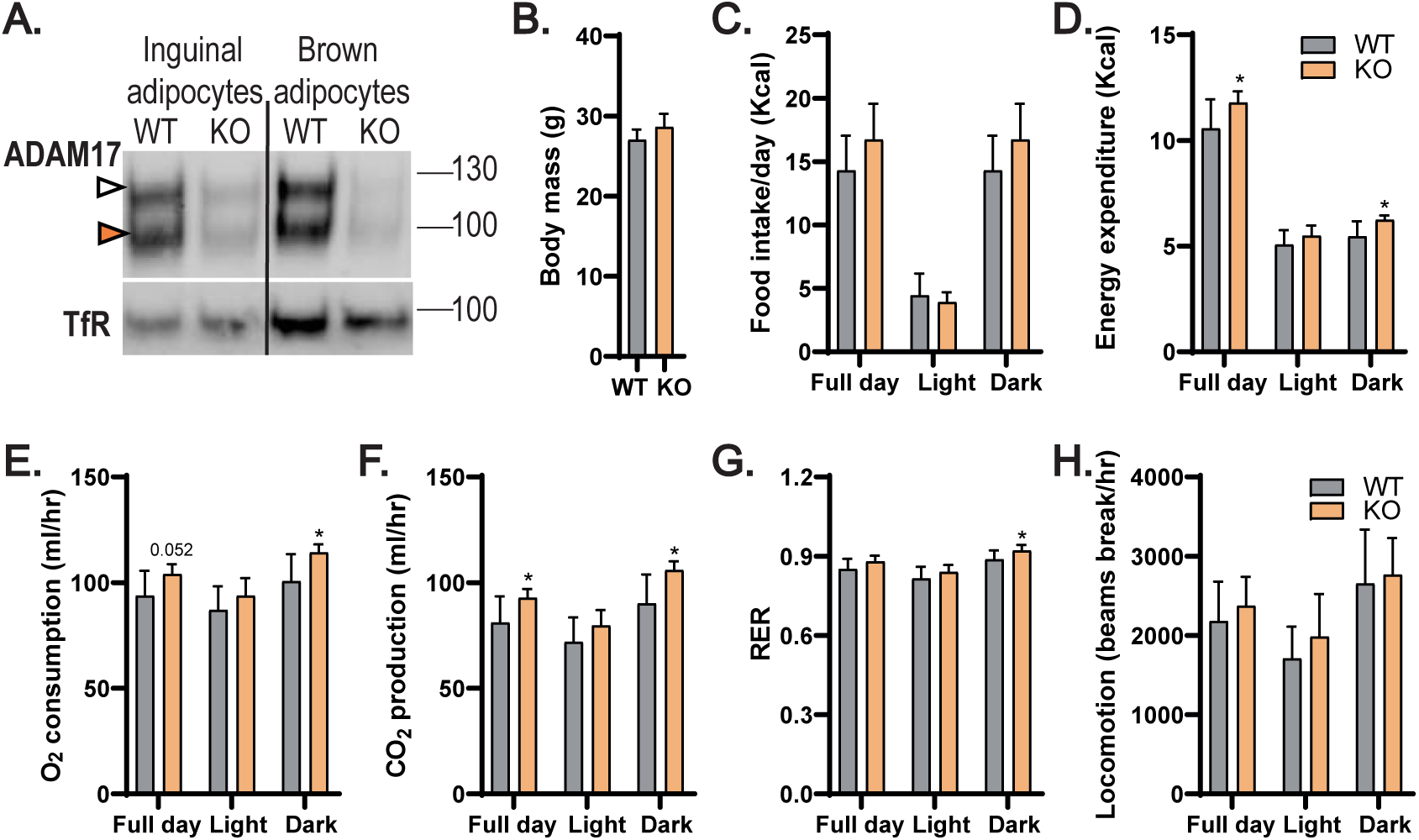
Lean mice null for ADAM17 in adipocytes have enhanced energy expenditure. **(A)**, Immunoblot of ADAM17 in adipocytes differentiated from WT and KO inguinal WAT and interscapular BAT. **(B-C)**, Body mass and 24 h food intake of lean WT and KO mice (WT n=8, KO n=7). **(D-G)**, Energy expenditure, oxygen consumption, carbon dioxide production, and respiratory exchange ratio of lean WT and KO mice (WT n=8, KO n=7). **(H)**, Locomotion of lean WT and KO mice (WT n=8, KO n=7). Results presented as mean ± SD. *P<0.05, **P<0.01, ***P<0.001, ****P<0.0001.

**Supplementary Figure 2:**
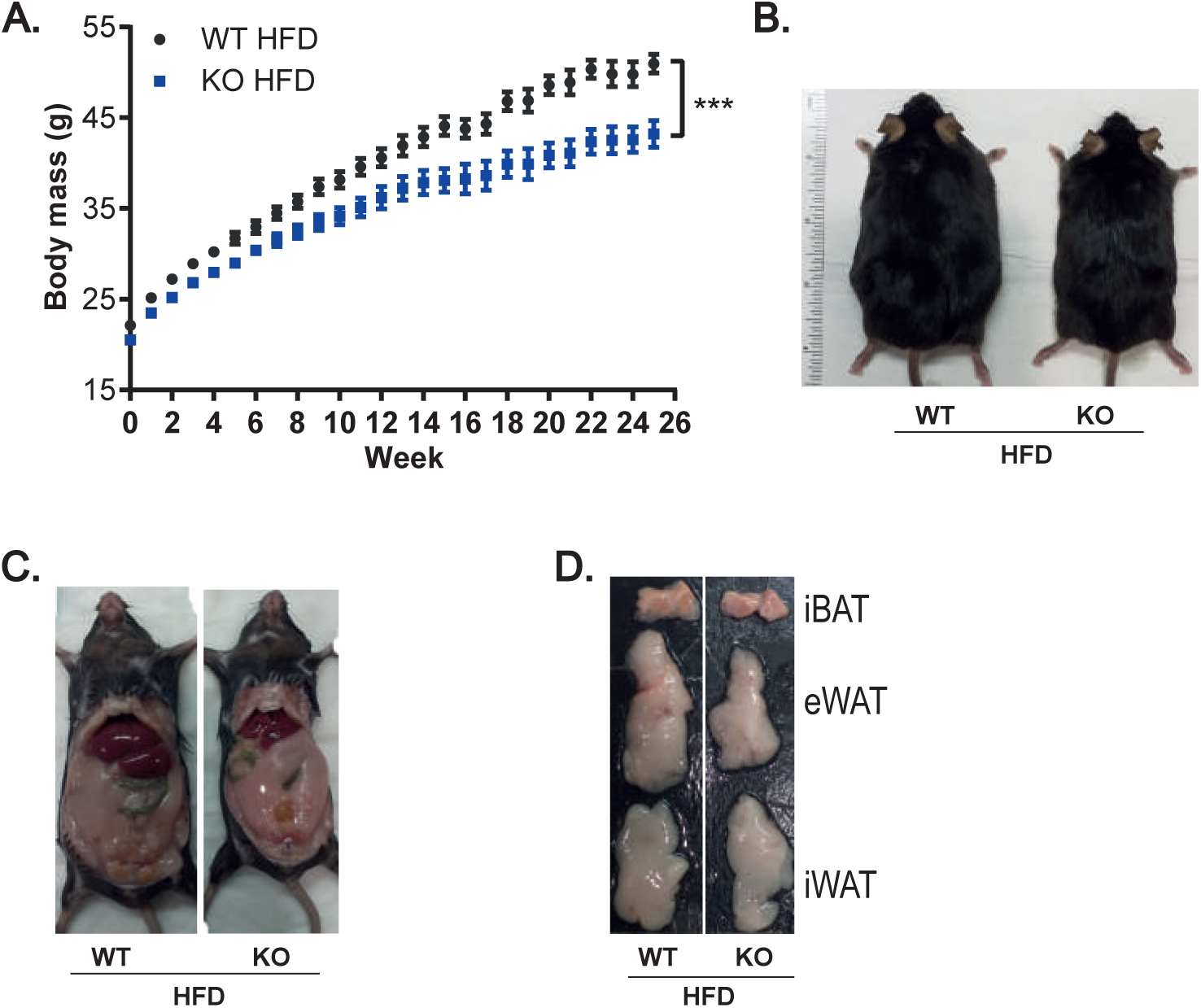
Mice null for ADAM17 in adipose tissue are less obese upon HFD feeding. (**A**), Change in body weight in WT and KO mice over the 26-week period of HFD feeding (WT n=20, KO n=18). **(B-D)**, Images of obese (B) WT and KO mice, of their abdominal organs (**C**), and of their interscapular BAT, inguinal WAT, and epididymal WAT (**D**) after 26 weeks of HFD feeding. Results presented as mean ± SD. *P<0.05, **P<0.01, ***P<0.001, ****P<0.0001.

**Supplementary Figure 3:**
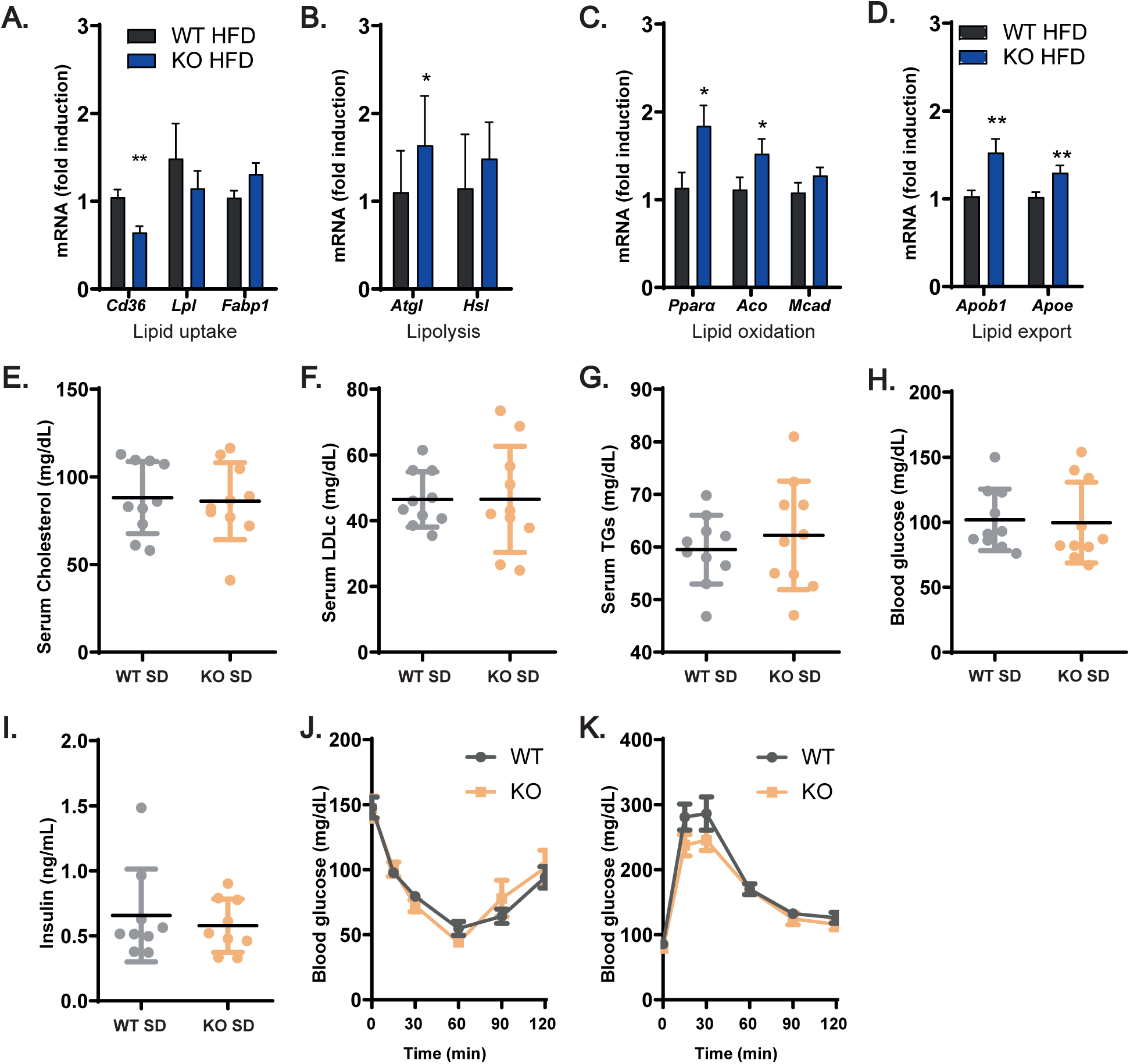
Loss of ADAM17 in adipocytes promotes lipid hepatic lipid catabolism in obesity but has no metabolic impact in lean condition. **(A-D)**, mRNA expression of genes involved in lipid uptake, lipolysis, lipid oxidation, and lipid export in the livers of obese WT and KO mice (WT n=13, KO n=12). **(E-G)**, Fasting serum levels of total cholesterol, LDLc, and triglycerides (TGs) in lean WT and KO mice (n=10). **(H-I)**, Fasting blood glucose and serum insulin levels of lean WT and KO mice. **(J-K),** Insulin and glucose tolerance tests in lean WT and KO mice (n=10). Results presented as mean ± SD. *P<0.05, **P<0.01, ***P<0.001, ****P<0.0001.

**Supplementary Figure 4:**
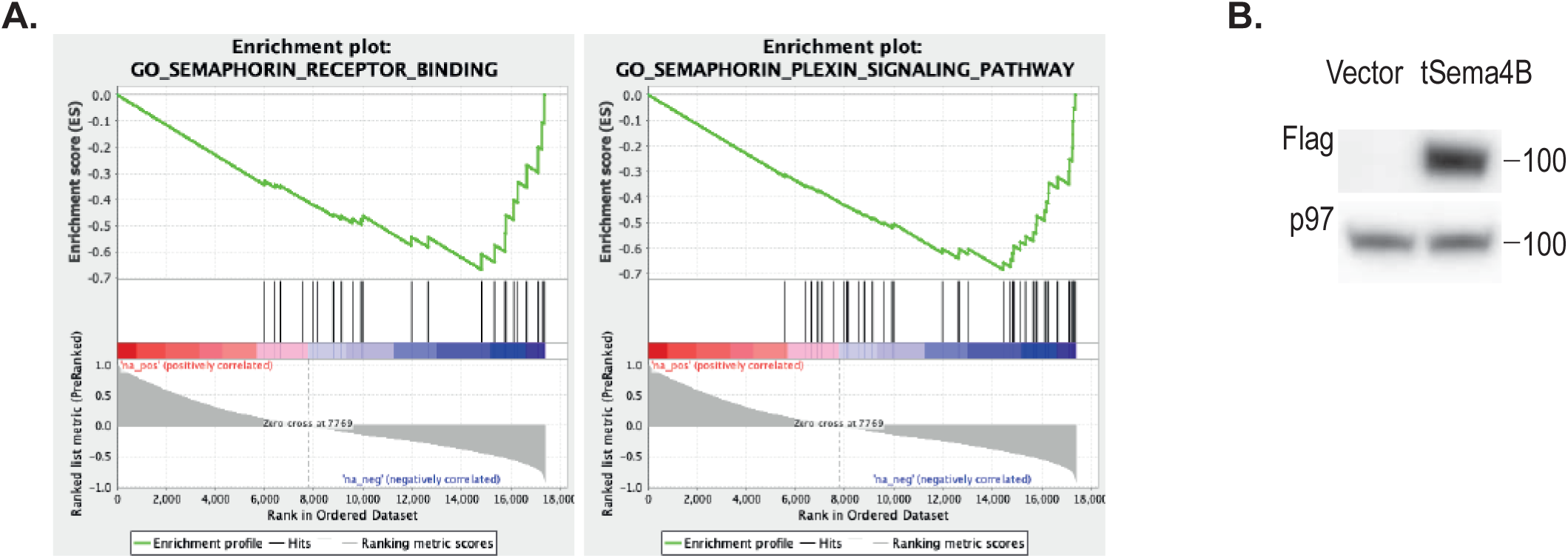
Semaphorin signaling inversely correlates with UCP1 expression. **(A)**, Gene set enrichment analysis (GSEA) from RNA seq data from Din et al. Genes enrich for the GO:TERM “semaphorin receptor binding” and “semaphorin plexin signaling pathway” negatively correlate with *UCP1* mRNA expression. **(B)**, Immunoblot for flag-tagged sema4B in lysates from immortalised primary brown adipocytes transduced with empty vector or truncated sema4B.

**Table 1.**
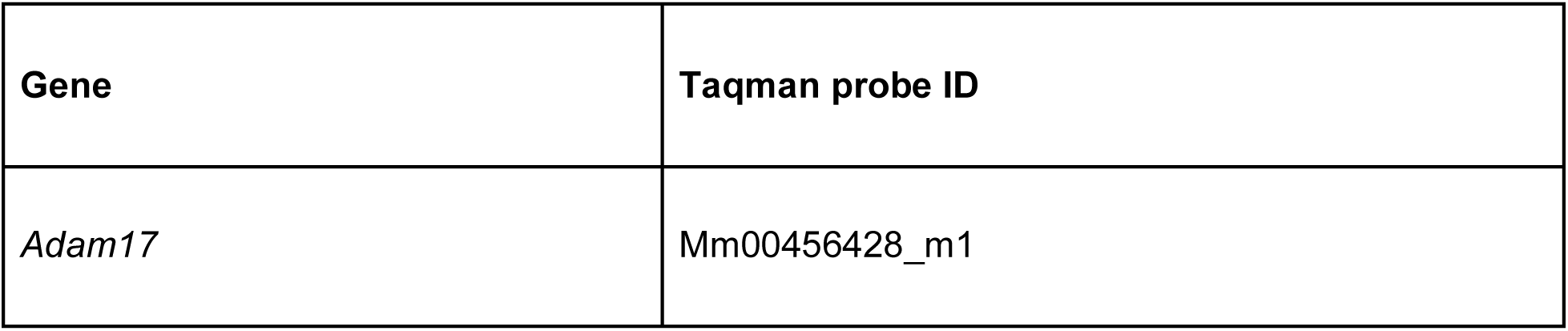
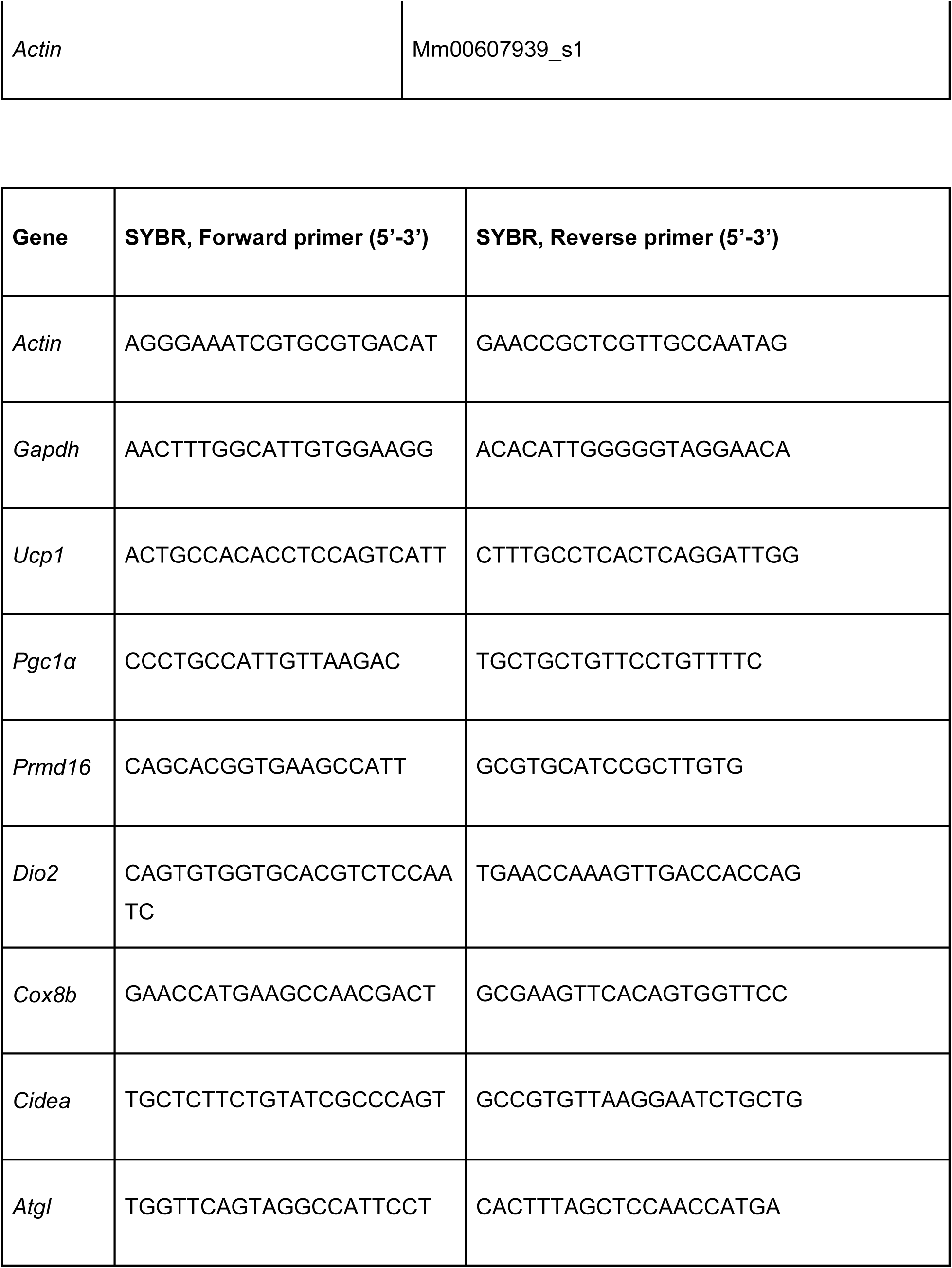
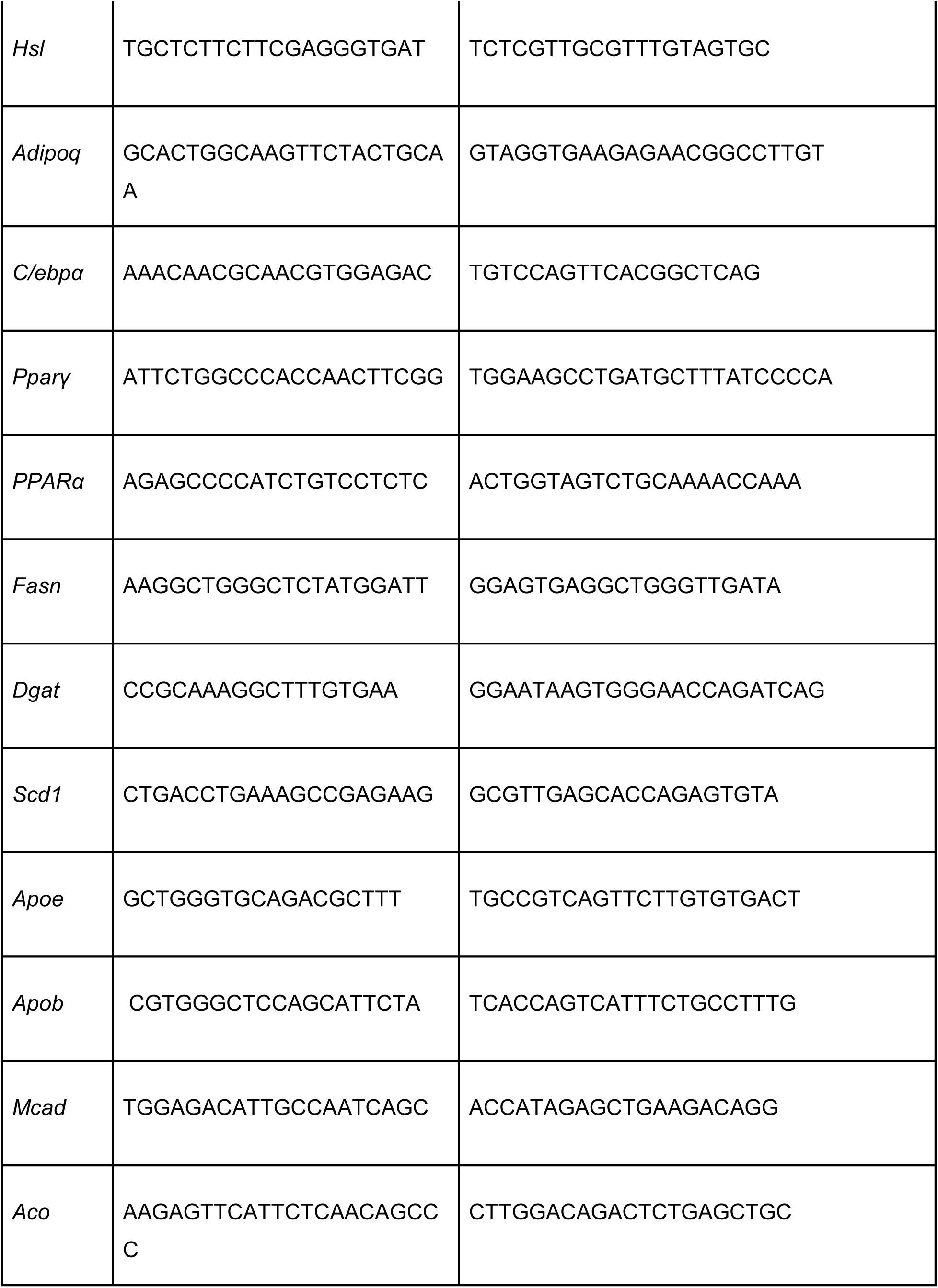
List of primers and Taqman probes

## References

[1] White, C. L., Purpera, M. N., Ballard, K., and Morrison, C. D. 2010. Decreased food intake following overfeeding involves leptin-dependent and leptin-independent mechanisms. Physiol Behav. 100, 4, 408–416.

[2] Ravussin, Y., Edwin, E., Gallop, M., Xu, L., Bartolomé, A., Kraakman, M. J., LeDuc, C. A., and Ferrante, A. W. 2018. Evidence for a Non-leptin System that Defends against Weight Gain in Overfeeding. Cell Metab. 28, 2, 289–299.e5.

[3] Richards, P., Thornberry, N. A., and Pinto, S. 2021. The gut-brain axis: Identifying new therapeutic approaches for type 2 diabetes, obesity, and related disorders. Mol Metab. 46, 101175.

[4] Müller, T. D., Blüher, M., Tschöp, M. H., and DiMarchi, R. D. 2021. Anti-obesity drug discovery: advances and challenges. Nature Reviews Drug Discovery. 1–23.

[5] Loos, R. J. F. and Yeo, G. S. H. 2022. The genetics of obesity: from discovery to biology. Nat Rev Genet. 23, 2, 120–133.

[6] Harvey, I., Boudreau, A., and Stephens, J. M. 2020. Adipose tissue in health and disease. Open Biol. 10, 12, 200291.

[7] Zwick, R. K., Guerrero-Juarez, C. F., Horsley, V., and Plikus, M. V. 2018. Anatomical, Physiological, and Functional Diversity of Adipose Tissue. Cell Metab. 27, 1, 68–83.

[8] Desruisseaux, M. S., Nagajyothi, Trujillo, M. E., Tanowitz, H. B., and Scherer, P. E. 2007. Adipocyte, adipose tissue, and infectious disease. Infect Immun. 75, 3, 1066–1078.

[9] Klaus, S. 2004. Adipose tissue as a regulator of energy balance. Curr Drug Targets. 5, 3, 241–250.

[10] Choe, S. S., Huh, J. Y., Hwang, I. J., Kim, J. I., and Kim, J. B. 2016. Adipose Tissue Remodeling: Its Role in Energy Metabolism and Metabolic Disorders. Front Endocrinol (Lausanne). 7, 30.

[11] Scheja, L. and Heeren, J. 2019. The endocrine function of adipose tissues in health and cardiometabolic disease. Nat Rev Endocrinol. 15, 9, 507–524.

[12] Rexford, A. and Flier, J. S. 2000. Leptin. Annu Rev Physiol. 62, 7, 413–437.

[13] Jequier, E. 2002. Leptin signaling, adiposity, and energy balance. Annals of the New York Academy of Sciences. 967, 1, 379–388.

[14] Griggio, M. A., Richard, D., and Leblanc, J. 1992. Effects of fasting and food restriction on sympathetic activity in brown adipose tissue in mice. J Comp Physiol B. 162, 7, 602–606.

[15] Kawate, R., Talan, M. I., and Engel, B. T. 1994. Sympathetic nervous activity to brown adipose tissue increases in cold-tolerant mice. Physiol Behav. 55, 5, 921–925.

[16] Cypess, A. M., Chen, Y. C., Sze, C., Wang, K., English, J., Chan, O., Holman, A. R., Tal, I., Palmer, M. R., Kolodny, G. M., and Kahn, C. R. 2012. Cold but not sympathomimetics activates human brown adipose tissue in vivo. Proc Natl Acad Sci U S A. 109, 25, 10001–10005.

[17] Bartness, T. J. and Song, C. K. 2007. Thematic review series: adipocyte biology. Sympathetic and sensory innervation of white adipose tissue. J Lipid Res. 48, 8, 1655–1672.

[18] François, M., Torres, H., Huesing, C., Zhang, R., Saurage, C., Lee, N., Qualls-Creekmore, E., Yu, S., Morrison, C. D., Burk, D., Berthoud, H. R., and Münzberg, H. 2019. Sympathetic innervation of the interscapular brown adipose tissue in mouse. Ann N Y Acad Sci. 1454, 1, 3–13.

[19] Münzberg, H., Floyd, E., and Chang, J. S. 2021. Sympathetic Innervation of White Adipose Tissue: to Beige or Not to Beige. Physiology (Bethesda). 36, 4, 246–255.

[20] Duncan, R. E., Ahmadian, M., Jaworski, K., Sarkadi-Nagy, E., and Sul, H. S. 2007. Regulation of lipolysis in adipocytes. Annu Rev Nutr. 27, 79–101.

[21] Grabner, G. F., Xie, H., Schweiger, M., and Zechner, R. 2021. Lipolysis: cellular mechanisms for lipid mobilization from fat stores. Nat Metab. 3, 11, 1445–1465.

[22] Collins, S. 2011. β-Adrenoceptor Signaling Networks in Adipocytes for Recruiting Stored Fat and Energy Expenditure. Front Endocrinol (Lausanne). 2, 102.

[23] Cannon, B. and Nedergaard, J. 2004. Brown adipose tissue: function and physiological significance. Physiol Rev. 84, 1, 277–359.

[24] Wade, G., McGahee, A., Ntambi, J. M., and Simcox, J. 2021. Lipid Transport in Brown Adipocyte Thermogenesis. Front Physiol. 12, 787535.

[25] Crichton, P. G., Lee, Y., and Kunji, E. R. 2017. The molecular features of uncoupling protein 1 support a conventional mitochondrial carrier-like mechanism. Biochimie. 134, 35–50.

[26] Fedorenko, A., Lishko, P. V., and Kirichok, Y. 2012. Mechanism of fatty-acid-dependent UCP1 uncoupling in brown fat mitochondria. Cell. 151, 2, 400–413.

[27] Wu, J., Boström, P., Sparks, L. M., Ye, L., Choi, J. H., Giang, A. H., Khandekar, M., Virtanen, K. A., Nuutila, P., Schaart, G., Huang, K., Tu, H., van Marken Lichtenbelt, W. D., Hoeks, J., Enerbäck, S., Schrauwen, P., and Spiegelman, B. M. 2012. Beige adipocytes are a distinct type of thermogenic fat cell in mouse and human. Cell. 150, 2, 366–376.

[28] Petrovic, N., Walden, T. B., and Shabalina…, I. G. 2010. … proliferator-activated receptor γ (PPARγ) activation of epididymally derived white adipocyte cultures reveals a population of thermogenically competent, UCP1 …. Journal of Biological ….

[29] Cannon, B. and Nedergaard, J. 2011. Nonshivering thermogenesis and its adequate measurement in metabolic studies. J Exp Biol. 214, Pt 2, 242–253.

[30] Roesler, A. and Kazak, L. 2020. UCP1-independent thermogenesis. Biochem J. 477, 3, 709–725.

[31] Singh, A. M., Zhang, L., Avery, J., Yin, A., Du, Y., Wang, H., Li, Z., Fu, H., Yin, H., and Dalton, S. 2020. Human beige adipocytes for drug discovery and cell therapy in metabolic diseases. Nat Commun. 11, 1, 2758.

[32] Thyagarajan, B. and Foster, M. T. 2017. Beiging of white adipose tissue as a therapeutic strategy for weight loss in humans. Horm Mol Biol Clin Investig. 31, 2, j/hmbci.2017.31.issue-2/hmbci.

[33] Moss, M. L., Jin, S.-L. C., Milla, M. E., Burkhart, W., Carter, H. L., Chen, W.-J., Clay, W. C., Didsbury, J. R., Hassler, D., and Hoffman, C. R. 1997. Cloning of a disintegrin metalloproteinase that processes precursor tumour-necrosis factor-α. Nature. 385, 6618, 733–736.

[34] Black, R. A., Rauch, C. T., Kozlosky, C. J., Peschon, J. J., Slack, J. L., Wolfson, M. F., Castner, B. J., Stocking, K. L., Reddy, P., and Srinivasan, S. 1997. A metalloproteinase disintegrin that releases tumour-necrosis factor-α from cells. Nature. 385, 6618, 729–733.

[35] Zunke, F. and Rose-John, S. 2017. The shedding protease ADAM17: Physiology and pathophysiology. Biochim Biophys Acta Mol Cell Res. 1864, 11 Pt B, 2059-2070.

[36] Schumacher, N. and Rose-John, S. 2019. ADAM17 Activity and IL-6 Trans-Signaling in Inflammation and Cancer. Cancers (Basel). 11, E1736.

[37] Lisi, S., D’Amore, M., and Sisto, M. 2014. ADAM17 at the interface between inflammation and autoimmunity. Immunol Lett. 162, 1 Pt A, 159-169.

[38] Ishii, S., Isozaki, T., Furuya, H., Takeuchi, H., Tsubokura, Y., Inagaki, K., and Kasama, T. 2018. ADAM-17 is expressed on rheumatoid arthritis fibroblast-like synoviocytes and regulates proinflammatory mediator expression and monocyte adhesion. Arthritis Res Ther. 20, 1, 159.

[39] Saad, M. I., Rose-John, S., and Jenkins, B. J. 2019. ADAM17: An Emerging Therapeutic Target for Lung Cancer. Cancers (Basel). 11, 9, E1218.

[40] Gelling, R. W., Yan, W., Al-Noori, S., Pardini, A., Morton, G. J., Ogimoto, K., Schwartz, M. W., and Dempsey, P. J. 2008. Deficiency of TNFalpha converting enzyme (TACE/ADAM17) causes a lean, hypermetabolic phenotype in mice. Endocrinology. 149, 12, 6053–6064.

[41] Serino, M., Menghini, R., Fiorentino, L., Amoruso, R., Mauriello, A., Lauro, D., Sbraccia, P., Hribal, M. L., Lauro, R., and Federici, M. 2007. Mice heterozygous for tumor necrosis factor-alpha converting enzyme are protected from obesity-induced insulin resistance and diabetes. Diabetes. 56, 10, 2541–2546.

[42] Hotamisligil, G. S., Shargill, N. S., and Spiegelman, B. M. 1993. Adipose expression of tumor necrosis factor-alpha: direct role in obesity-linked insulin resistance. Science. 259, 5091, 87–91.

[43] Hotamisligil, G. S. 1999. The role of TNFα and TNF receptors in obesity and insulin resistance. Journal of internal medicine. 245, 6, 621–625.

[44] Uysal, K. T., Wiesbrock, S. M., Marino, M. W., and Hotamisligil, G. S. 1997. Protection from obesity-induced insulin resistance in mice lacking TNF-α function. Nature. 389, 6651, 610–614.

[45] Badenes, M., Amin, A., González-García, I., Félix, I., Burbridge, E., Cavadas, M., Ortega, F. J., de Carvalho, É., Faísca, P., Carobbio, S., Seixas, E., Pedroso, D., Neves-Costa, A., Moita, L. F., Fernández-Real, J. M., Vidal-Puig, A., Domingos, A., López, M., and Adrain, C. 2020. Deletion of iRhom2 protects against diet-induced obesity by increasing thermogenesis. Mol Metab. 31, 67–84.

[46] Adrain, C., Zettl, M., Christova, Y., Taylor, N., and Freeman, M. 2012. Tumor necrosis factor signaling requires iRhom2 to promote trafficking and activation of TACE. Science. 335, 6065, 225–228.

[47] McIlwain, D. R., Lang, P. A., Maretzky, T., Hamada, K., Ohishi, K., Maney, S. K., Berger, T., Murthy, A., Duncan, G., Xu, H. C., Lang, K. S., Häussinger, D., Wakeham, A., Itie-Youten, A., Khokha, R., Ohashi, P. S., Blobel, C. P., and Mak, T. W. 2012. iRhom2 regulation of TACE controls TNF-mediated protection against Listeria and responses to LPS. Science. 335, 6065, 229–232.

[48] Maretzky, T., McIlwain, D. R., Issuree, P. D., Li, X., Malapeira, J., Amin, S., Lang, P. A., Mak, T. W., and Blobel, C. P. 2013. iRhom2 controls the substrate selectivity of stimulated ADAM17-dependent ectodomain shedding. Proc Natl Acad Sci U S A. 110, 28, 11433–11438.

[49] Cavadas, M., Oikonomidi, I., Gaspar, C. J., Burbridge, E., Badenes, M., Félix, I., Bolado, A., Hu, T., Bileck, A., Gerner, C., Domingos, P. M., von Kriegsheim, A., and Adrain, C. 2017. Phosphorylation of iRhom2 Controls Stimulated Proteolytic Shedding by the Metalloprotease ADAM17/TACE. Cell Rep. 21, 3, 745–757.

[50] Grieve, A. G., Xu, H., Künzel, U., Bambrough, P., Sieber, B., and Freeman, M. 2017. Phosphorylation of iRhom2 at the plasma membrane controls mammalian TACE-dependent inflammatory and growth factor signalling. Elife. 6, e23968.

[51] Oikonomidi, I., Burbridge, E., Cavadas, M., Sullivan, G., Collis, B., Naegele, H., Clancy, D., Brezinova, J., Hu, T., Bileck, A., Gerner, C., Bolado, A., von Kriegsheim, A., Martin, S. J., Steinberg, F., Strisovsky, K., and Adrain, C. 2018. iTAP, a novel iRhom interactor, controls TNF secretion by policing the stability of iRhom/TACE. Elife. 7, e35032.

[52] Künzel, U., Grieve, A. G., Meng, Y., Sieber, B., Cowley, S. A., and Freeman, M. 2018. FRMD8 promotes inflammatory and growth factor signalling by stabilising the iRhom/ADAM17 sheddase complex. Elife. 7, e35012.

[53] Badenes, M., Burbridge, E., Oikonomidi, I., Amin, A., de Carvalho, É., Kosack, L., Domingos, P., Faísca, P., and Adrain, C. 2022. The ADAM17 sheddase complex regulator iTAP modulates inflammation, epithelial repair, and tumor growth.

[54] Westerterp, K. R. 2004. Diet induced thermogenesis. Nutr Metab (Lond). 1, 5.

[55] Saito, M., Matsushita, M., Yoneshiro, T., and Okamatsu-Ogura, Y. 2020. Brown Adipose Tissue, Diet-Induced Thermogenesis, and Thermogenic Food Ingredients: From Mice to Men. Front Endocrinol (Lausanne). 11, 222.

[56] Fischer, A. W., Schlein, C., Cannon, B., Heeren, J., and Nedergaard, J. 2019. Intact innervation is essential for diet-induced recruitment of brown adipose tissue. Am J Physiol Endocrinol Metab. 316, 3, E487–E503.

[57] Prenzel, N., Zwick, E., Daub, H., Leserer, M., Abraham, R., Wallasch, C., and Ullrich, A. 1999. EGF receptor transactivation by G-protein-coupled receptors requires metalloproteinase cleavage of proHB-EGF. Nature. 402, 6764, 884–888.

[58] Gooz, M., Gooz, P., Luttrell, L. M., and Raymond, J. R. 2006. 5-HT2A receptor induces ERK phosphorylation and proliferation through ADAM-17 tumor necrosis factor-α-converting enzyme (TACE) activation and heparin …. Journal of Biological Chemistry.

[59] Yin, J. and Yu, F. S. 2009. ERK1/2 mediate wounding- and G-protein-coupled receptor ligands-induced EGFR activation via regulating ADAM17 and HB-EGF shedding. Invest Ophthalmol Vis Sci. 50, 1, 132–139.

[60] Horiuchi, K., Kimura, T., Miyamoto, T., Takaishi, H., Okada, Y., Toyama, Y., and Blobel, C. P. 2007. Cutting edge: TNF-alpha-converting enzyme (TACE/ADAM17) inactivation in mouse myeloid cells prevents lethality from endotoxin shock. J Immunol. 179, 5, 2686–2689.

[61] Eguchi, J., Wang, X., Yu, S., Kershaw, E. E., Chiu, P. C., Dushay, J., Estall, J. L., Klein, U., Maratos-Flier, E., and Rosen, E. D. 2011. Transcriptional control of adipose lipid handling by IRF4. Cell Metab. 13, 3, 249–259.

[62] Carobbio, S., Guenantin, A. C., Bahri, M., Rodriguez-Fdez, S., Honig, F., Kamzolas, I., Samuelson, I., Long, K., Awad, S., Lukovic, D., Erceg, S., Bassett, A., Mendjan, S., Vallier, L., Rosen, B. S., Chiarugi, D., and Vidal-Puig, A. 2021. Unraveling the Developmental Roadmap toward Human Brown AdiposeTissue. Stem Cell Reports. 16, 4, 1010.

[63] Tran, K. V., Brown, E. L., DeSouza, T., Jespersen, N. Z., Nandrup-Bus, C., Yang, Q., Yang, Z., Desai, A., Min, S. Y., Rojas-Rodriguez, R., Lundh, M., Feizi, A., Willenbrock, H., Larsen, T. J., Severinsen, M. C. K., Malka, K., Mozzicato, A. M., Deshmukh, A. S., Emanuelli, B., Pedersen, B. K., Fitzgibbons, T., Scheele, C., Corvera, S., and Nielsen, S. 2020. Human thermogenic adipocyte regulation by the long noncoding RNA LINC00473. Nat Metab. 2, 5, 397–412.

[64] Tüshaus, J., Müller, S. A., Kataka, E. S., Zaucha, J., Sebastian Monasor, L., Su, M., Güner, G., Jocher, G., Tahirovic, S., Frishman, D., Simons, M., and Lichtenthaler, S. F. 2020. An optimized quantitative proteomics method establishes the cell type-resolved mouse brain secretome. EMBO J. 39, 20, e105693.

[65] Tüshaus, J., Müller, S. A., Shrouder, J., Arends, M., Simons, M., Plesnila, N., Blobel, C. P., and Lichtenthaler, S. F. 2021. The pseudoprotease iRhom1 controls ectodomain shedding of membrane proteins in the nervous system. FASEB J. 35, 11, e21962.

[66] Burkhardt, C., Müller, M., Badde, A., Garner, C. C., Gundelfinger, E. D., and Püschel, A. W. 2005. Semaphorin 4B interacts with the post-synaptic density protein PSD-95/SAP90 and is recruited to synapses through a C-terminal PDZ-binding motif. FEBS Lett. 579, 17, 3821–3828.

[67] Adrain, C., Strisovsky, K., Zettl, M., Hu, L., Lemberg, M. K., and Freeman, M. 2011. Mammalian EGF receptor activation by the rhomboid protease RHBDL2. EMBO Rep. 12, 5, 421–427.

[68] Naviaux, R. K., Costanzi, E., Haas, M., and Verma, I. M. 1996. The pCL vector system: rapid production of helper-free, high-titer, recombinant retroviruses. J Virol. 70, 8, 5701–5705.

[69] Cavadas, M., Oikonomidi, I., Gaspar, C. J., Burbridge, E., Badenes, M., Félix, I., Bolado, A., Hu, T., Bileck, A., Gerner, C., Domingos, P. M., von Kriegsheim, A., and Adrain, C. 2017. Phosphorylation of iRhom2 Controls Stimulated Proteolytic Shedding by the Metalloprotease ADAM17/TACE. Cell Rep. 21, 3, 745–757.

[70] Klein, J., Fasshauer, M., Ito, M., Lowell, B. B., Benito, M., and Kahn, C. R. 1999. β3-Adrenergic stimulation differentially inhibits insulin signaling and decreases insulin-induced glucose uptake in brown adipocytes. Journal of Biological Chemistry. 274, 49, 34795–34802.

[71] Picelli, S., Faridani, O. R., Björklund, A. K., Winberg, G., Sagasser, S., and Sandberg, R. 2014. Full-length RNA-seq from single cells using Smart-seq2. Nat Protoc. 9, 1, 171–181.

[72] Baym, M., Kryazhimskiy, S., Lieberman, T. D., Chung, H., Desai, M. M., and Kishony, R. 2015. Inexpensive multiplexed library preparation for megabase-sized genomes. PLoS One. 10, 5, e0128036.

[73] Dobin, A., Davis, C. A., Schlesinger, F., Drenkow, J., Zaleski, C., Jha, S., Batut, P., Chaisson, M., and Gingeras, T. R. 2013. STAR: ultrafast universal RNA-seq aligner. Bioinformatics. 29, 1, 15–21.

[74] Anders, S., Pyl, P. T., and Huber, W. 2015. HTSeq--a Python framework to work with high-throughput sequencing data. Bioinformatics. 31, 2, 166–169.

[75] Love, M. I., Huber, W., and Anders, S. 2014. Moderated estimation of fold change and dispersion for RNA-seq data with DESeq2. Genome Biol. 15, 12, 550.

[76] Sherman, B. T., Hao, M., Qiu, J., Jiao, X., Baseler, M. W., Lane, H. C., Imamichi, T., and Chang, W. 2022. DAVID: a web server for functional enrichment analysis and functional annotation of gene lists (2021 update). Nucleic Acids Res. gkac194.

[77] Prenzel, N., Zwick, E., Daub, H., Leserer, M., Abraham, R., Wallasch, C., and Ullrich, A. 1999. EGF receptor transactivation by G-protein-coupled receptors requires metalloproteinase cleavage of proHB-EGF. Nature. 402, 6764, 884–888.

[78] Palanisamy, S., Xue, C., Ishiyama, S., Naga Prasad, S. V., and Gabrielson, K. 2021. GPCR-ErbB transactivation pathways and clinical implications. Cell Signal. 86, 110092.

[79] Collins, S. 2022. β-Adrenergic Receptors and Adipose Tissue Metabolism: Evolution of an Old Story. Annual Review of Physiology. 84, 1–16.

[80] Shook, R. P., Hand, G. A., Paluch, A. E., Wang, X., Moran, R., Hébert, J. R., Jakicic, J. M., and Blair, S. N. 2016. High respiratory quotient is associated with increases in body weight and fat mass in young adults. Eur J Clin Nutr. 70, 10, 1197–1202.

[81] Burke, S. J., Batdorf, H. M., Martin, T. M., Burk, D. H., Noland, R. C., Cooley, C. R., Karlstad, M. D., Johnson, W. D., and Collier, J. J. 2018. Liquid Sucrose Consumption Promotes Obesity and Impairs Glucose Tolerance Without Altering Circulating Insulin Levels. Obesity (Silver Spring). 26, 7, 1188–1196.

[82] Jo, J., Gavrilova, O., Pack, S., Jou, W., Mullen, S., Sumner, A. E., Cushman, S. W., and Periwal, V. 2009. Hypertrophy and/or Hyperplasia: Dynamics of Adipose Tissue Growth. PLoS Comput Biol. 5, 3, e1000324.

[83] Attie, A. D. and Scherer, P. E. 2009. Adipocyte metabolism and obesity. J Lipid Res. 50 Suppl, S395–9.

[84] Geisler, C. E. and Renquist, B. J. 2017. Hepatic lipid accumulation: cause and consequence of dysregulated glucoregulatory hormones. J Endocrinol. 234, 1, R1–R21.

[85] Koonen, D. P., Jacobs, R. L., Febbraio, M., Young, M. E., Soltys, C. L., Ong, H., Vance, D. E., and Dyck, J. R. 2007. Increased hepatic CD36 expression contributes to dyslipidemia associated with diet-induced obesity. Diabetes. 56, 12, 2863–2871.

[86] De Taeye, B. M., Novitskaya, T., McGuinness, O. P., Gleaves, L., Medda, M., Covington, J. W., and Vaughan, D. E. 2007. Macrophage TNF-alpha contributes to insulin resistance and hepatic steatosis in diet-induced obesity. Am J Physiol Endocrinol Metab. 293, 3, E713–25.

[87] Sugawara, K., Schneider, M. R., Dahlhoff, M., Kloepper, J. E., and Paus, R. 2010. Cutaneous consequences of inhibiting EGF receptor signaling in vivo: normal hair follicle development, but retarded hair cycle induction and inhibition of adipocyte growth in Egfr(Wa5) mice. J Dermatol Sci. 57, 3, 155–161.

[88] Choung, S., Kim, J. M., Joung, K. H., Lee, E. S., Kim, H. J., and Ku, B. J. 2019. Epidermal growth factor receptor inhibition attenuates non-alcoholic fatty liver disease in diet-induced obese mice. PloS one. 14, 2, e0210828.

[89] Timper, K., Denson, J. L., Steculorum, S. M., Heilinger, C., Engström-Ruud, L., Wunderlich, C. M., Rose-John, S., Wunderlich, F. T., and Brüning, J. C. 2017. IL-6 Improves Energy and Glucose Homeostasis in Obesity via Enhanced Central IL-6 trans-Signaling. Cell Rep. 19, 2, 267–280.

[90] Schöbitz, B., Pezeshki, G., Pohl, T., Hemmann, U., Heinrich, P. C., Holsboer, F., and Reul, J. M. H. M. 1995. Soluble interleukin-6 (IL-6) receptor augments central effects of IL- 6 in vivo. The FASEB Journal. 9, 8, 659–664.

[91] Wallenius, V., Wallenius, K., Ahrén, B., Rudling, M., Carlsten, H., Dickson, S. L., Ohlsson, C., and Jansson, J.-O. 2002. Interleukin-6-deficient mice develop mature-onset obesity. Nature medicine. 8, 1, 75–79.

[92] Wallenius, K., Wallenius, V., Sunter, D., Dickson, S. L., and Jansson, J.-O. 2002. Intracerebroventricular interleukin-6 treatment decreases body fat in rats. Biochemical and biophysical research communications. 293, 1, 560–565.

[93] Tüshaus, J., Müller, S. A., Kataka, E. S., Zaucha, J., Sebastian Monasor, L., Su, M., Güner, G., Jocher, G., Tahirovic, S., Frishman, D., Simons, M., and Lichtenthaler, S. F. 2020. An optimized quantitative proteomics method establishes the cell type-resolved mouse brain secretome. EMBO J. 39, 20, e105693.

[94] Alto, L. T. and Terman, J. R. 2017. Semaphorins and their signaling mechanisms. Semaphorin signaling. 1–25.

[95] Ramseyer, V. D. and Granneman, J. G. 2016. Adrenergic regulation of cellular plasticity in brown, beige/brite and white adipose tissues. Adipocyte. 5, 2, 119–129.

[96] Christie, S. M., Hao, J., Tracy, E., Buck, M., Yu, J. S., and Smith, A. W. 2021. Interactions between semaphorins and plexin-neuropilin receptor complexes in the membranes of live cells. J Biol Chem. 297, 2, 100965.

[97] U Din, M., Saari, T., Raiko, J., Kudomi, N., Maurer, S. F., Lahesmaa, M., Fromme, T., Amri, E. Z., Klingenspor, M., Solin, O., Nuutila, P., and Virtanen, K. A. 2018. Postprandial Oxidative Metabolism of Human Brown Fat Indicates Thermogenesis. Cell Metab. 28, 2, 207–216.e3.

[98] Lorenzen, I., Lokau, J., Korpys, Y., Oldefest, M., Flynn, C. M., Künzel, U., Garbers, C., Freeman, M., Grötzinger, J., and Düsterhöft, S. 2016. Control of ADAM17 activity by regulation of its cellular localisation. Sci Rep. 6, 35067.

[99] Wang, Z. 2016. Transactivation of Epidermal Growth Factor Receptor by G Protein-Coupled Receptors: Recent Progress, Challenges and Future Research. Int J Mol Sci. 17, 1, E95.

[100] Fischer, A. W., Cannon, B., and Nedergaard, J. 2018. Optimal housing temperatures for mice to mimic the thermal environment of humans: An experimental study. Mol Metab. 7, 161–170.

[101] Škop, V., Guo, J., Liu, N., Xiao, C., Hall, K. D., Gavrilova, O., and Reitman, M. L. 2020. Mouse Thermoregulation: Introducing the Concept of the Thermoneutral Point. Cell Rep. 31, 2, 107501.

[102] Ravussin, Y., LeDuc, C. A., Watanabe, K., and Leibel, R. L. 2012. Effects of ambient temperature on adaptive thermogenesis during maintenance of reduced body weight in mice. Am J Physiol Regul Integr Comp Physiol. 303, 4, R438–48.

[103] Yang, T. and Terman, J. R. 2012. 14-3-3ε couples protein kinase A to semaphorin signaling and silences plexin RasGAP-mediated axonal repulsion. Neuron. 74, 1, 108–121.

[104] Kumanogoh, A., Watanabe, C., Lee, I., Wang, X., Shi, W., Araki, H., Hirata, H., Iwahori, K., Uchida, J., and Yasui, T. 2000. Identification of CD72 as a lymphocyte receptor for the class IV semaphorin CD100: a novel mechanism for regulating B cell signaling. Immunity. 13, 5, 621–631.

[105] Cho, J. Y., Chak, K., Andreone, B. J., Wooley, J. R., and Kolodkin, A. L. 2012. The extracellular matrix proteoglycan perlecan facilitates transmembrane semaphorin-mediated repulsive guidance. Genes Dev. 26, 19, 2222–2235.

[106] Kumanogoh, A., Marukawa, S., Suzuki, K., Takegahara, N., Watanabe, C., Ch’ng, E., Ishida, I., Fujimura, H., Sakoda, S., Yoshida, K., and Kikutani, H. 2002. Class IV semaphorin Sema4A enhances T-cell activation and interacts with Tim-2. Nature. 419, 6907, 629–633.

[107] Pasterkamp, R. J., Peschon, J. J., and Spriggs…, M. K. 2003. Semaphorin 7A promotes axon outgrowth through integrins and MAPKs. Nature.

[108] Nedergaard, J., Petrovic, N., Lindgren, E. M., Jacobsson, A., and Cannon, B. 2005. PPARgamma in the control of brown adipocyte differentiation. Biochim Biophys Acta. 1740, 2, 293–304.

[109] Manchado, C., Yubero, P., Viñas, O., Iglesias, R., Villarroya, F., Mampel, T., and Giralt, M. 1994. CCAAT/enhancer-binding proteins α and β in brown adipose tissue: evidence for a tissue-specific pattern of expression during development. Biochemical Journal. 302, 3, 695–700.

[110] Bos, J. L., Rehmann, H., and Wittinghofer, A. 2007. GEFs and GAPs: critical elements in the control of small G proteins. Cell. 129, 5, 865–877.

[111] Wang, Y., He, H., Srivastava, N., Vikarunnessa, S., Chen, Y. B., Jiang, J., Cowan, C. W., and Zhang, X. 2012. Plexins are GTPase-activating proteins for Rap and are activated by induced dimerization. Sci Signal. 5, 207, ra6.

[112] Biernacka, A., Dobaczewski, M., and Frangogiannis, N. G. 2011. TGF-β signaling in fibrosis. Growth Factors. 29, 5, 196–202.

[113] Kefaloyianni, E., Muthu, M. L., Kaeppler, J., Sun, X., Sabbisetti, V., Chalaris, A., Rose-John, S., Wong, E., Sagi, I., Waikar, S. S., Rennke, H., Humphreys, B. D., Bonventre, J. V., and Herrlich, A. 2016. ADAM17 substrate release in proximal tubule drives kidney fibrosis. JCI Insight. 1, 13, 87023.

[114] Matsui, Y., Tomaru, U., Miyoshi, A., Ito, T., Fukaya, S., Miyoshi, H., Atsumi, T., and Ishizu, A. 2014. Overexpression of TNF-α converting enzyme promotes adipose tissue inflammation and fibrosis induced by high fat diet. Exp Mol Pathol. 97, 3, 354–358.

[115] Wang, X., Oka, T., Chow, F. L., Cooper, S. B., Odenbach, J., Lopaschuk, G. D., Kassiri, Z., and Fernandez-Patron, C. 2009. Tumor necrosis factor-alpha-converting enzyme is a key regulator of agonist-induced cardiac hypertrophy and fibrosis. Hypertension. 54, 3, 575–582.

[116] Carvalheiro, T., Affandi, A. J., Malvar-Fernández, B., Dullemond, I., Cossu, M., Ottria, A., Mertens, J. S., Giovannone, B., Bonte-Mineur, F., Kok, M. R., Marut, W., Reedquist, K. A., Radstake, T. R., and García, S. 2019. Induction of Inflammation and Fibrosis by Semaphorin 4A in Systemic Sclerosis. Arthritis Rheumatol. 71, 10, 1711–1722.

[117] Jeon, K. I., Nehrke, K., and Huxlin, K. R. 2020. Semaphorin 3A potentiates the profibrotic effects of transforming growth factor-β1 in the cornea. Biochem Biophys Res Commun. 521, 2, 333–339.

[118] Peng, H. Y., Gao, W., Chong, F. R., Liu, H. Y., and Zhang, J. I. 2015. Semaphorin 4A enhances lung fibrosis through activation of Akt via PlexinD1 receptor. J Biosci. 40, 5, 855–862.

[119] Sang, Y., Tsuji, K., Fukushima, K., Takahashi, K., Kitamura, S., and Wada, J. 2021. Semaporin3A inhibitor ameliorates renal fibrosis through the regulation of JNK signaling. Am J Physiol Renal Physiol. 321, 6, F740–F756.

[120] Ruiz-Ojeda, F. J., Méndez-Gutiérrez, A., Aguilera, C. M., and Plaza-Díaz, J. 2019. Extracellular Matrix Remodeling of Adipose Tissue in Obesity and Metabolic Diseases. Int J Mol Sci. 20, 19, E4888.

[121] Mejhert, N., Wilfling, F., Esteve, D., Galitzky, J., Pellegrinelli, V., Kolditz, C. I., Viguerie, N., Tordjman, J., Näslund, E., Trayhurn, P., Lacasa, D., Dahlman, I., Stich, V., Lång, P., Langin, D., Bouloumié, A., Clément, K., and Rydén, M. 2013. Semaphorin 3C is a novel adipokine linked to extracellular matrix composition. Diabetologia. 56, 8, 1792–1801.

[122] Hasegawa, Y., Ikeda, K., Chen, Y., Alba, D. L., Stifler, D., Shinoda, K., Hosono, T., Maretich, P., Yang, Y., Ishigaki, Y., Chi, J., Cohen, P., Koliwad, S. K., and Kajimura, S. 2018. Repression of Adipose Tissue Fibrosis through a PRDM16-GTF2IRD1 Complex Improves Systemic Glucose Homeostasis. Cell Metab. 27, 1, 180–194.e6.

[123] Chouchani, E. T. and Kajimura, S. 2019. Metabolic adaptation and maladaptation in adipose tissue. Nat Metab. 1, 2, 189–200.

[124] Gonzalez Porras, M. A., Stojkova, K., Vaicik, M. K., Pelowe, A., Goddi, A., Carmona, A., Long, B., Qutub, A. A., Gonzalez, A., Cohen, R. N., and Brey, E. M. 2021. Integrins and extracellular matrix proteins modulate adipocyte thermogenic capacity. Sci Rep. 11, 1, 5442.

[125] van der Klaauw, A. A., Croizier, S., Mendes de Oliveira, E., Stadler, L. K. J., Park, S., Kong, Y., Banton, M. C., Tandon, P., Hendricks, A. E., Keogh, J. M., Riley, S. E., Papadia, S., Henning, E., Bounds, R., Bochukova, E. G., Mistry, V., O’Rahilly, S., Simerly, R. B., INTERVAL, UK10K, C., Minchin, J. E. N., Barroso, I., Jones, E. Y., Bouret, S. G., and Farooqi, I. S. 2019. Human Semaphorin 3 Variants Link Melanocortin Circuit Development and Energy Balance. Cell. 176, 4, 729-742.e18.

[126] Giordano, A., Coppari, R., Castellucci, M., and Cinti, S. 2001. Sema3a is produced by brown adipocytes and its secretion is reduced following cold acclimation. J Neurocytol. 30, 1, 5–10.

[127] Giordano, A., Cesari, P., Capparuccia, L., Castellucci, M., and Cinti, S. 2003. Sema3A and neuropilin-1 expression and distribution in rat white adipose tissue. J Neurocytol. 32, 4, 345–352.

[128] Lu, Q. and Zhu, L. 2020. The Role of Semaphorins in Metabolic Disorders. Int J Mol Sci. 21, 16, E5641.

[129] Liu, M., Xie, S., Liu, W., Li, J., Li, C., Huang, W., Li, H., Song, J., and Zhang, H. 2020. Mechanism of SEMA3G knockdown-mediated attenuation of high-fat diet-induced obesity. J Endocrinol. 244, 1, 223–236.

[130] Fong, K. P., Barry, C., Tran, A. N., Traxler, E. A., Wannemacher, K. M., Tang, H. Y., Speicher, K. D., Blair, I. A., Speicher, D. W., Grosser, T., and Brass, L. F. 2011. Deciphering the human platelet sheddome. Blood. 117, 1, e15–26.

[131] Motani, K. and Kosako, H. 2018. Activation of stimulator of interferon genes (STING) induces ADAM17-mediated shedding of the immune semaphorin SEMA4D. J Biol Chem. 293, 20, 7717–7726.

[132] Browne, K., Wang, W., Liu, R. Q., Piva, M., and O’Connor, T. P. 2012. Transmembrane semaphorin5B is proteolytically processed into a repulsive neural guidance cue. J Neurochem. 123, 1, 135–146.

[133] Romi, E., Gokhman, I., Wong, E., Antonovsky, N., Ludwig, A., Sagi, I., Saftig, P., Tessier-Lavigne, M., and Yaron, A. 2014. ADAM metalloproteases promote a developmental switch in responsiveness to the axonal repellant Sema3A. Nat Commun. 5, 4058.

[134] Ito, T., Bai, T., Tanaka, T., Yoshida, K., Ueyama, T., Miyajima, M., Negishi, T., Kawasaki, T., Takamatsu, H., Kikutani, H., Kumanogoh, A., and Yukawa, K. 2014. Estrogen-dependent proteolytic cleavage of semaphorin 4D and plexin-B1 enhances semaphorin 4D-induced apoptosis during postnatal vaginal remodeling in pubescent mice. PLoS One. 9, 5, e97909.

[135] Esselens, C., Malapeira, J., Colomé, N., Casal, C., Rodríguez-Manzaneque, J. C., Canals, F., and Arribas, J. 2010. The cleavage of semaphorin 3C induced by ADAMTS1 promotes cell migration. J Biol Chem. 285, 4, 2463–2473.

[136] Murumkar, P. R., Ghuge, R. B., Chauhan, M., Barot, R. R., Sorathiya, S., Choudhary, K. M., Joshi, K. D., and Yadav, M. R. 2020. Recent developments and strategies for the discovery of TACE inhibitors. Expert Opin Drug Discov. 15, 7, 779–801.

[137] Calligaris, M., Cuffaro, D., Bonelli, S., Spanò, D. P., Rossello, A., Nuti, E., and Scilabra, S. D. 2021. Strategies to Target ADAM17 in Disease: From its Discovery to the iRhom Revolution. Molecules. 26, 4, 944.

